# Distinguishing between- from within-site phase-amplitude coupling using antisymmetrized bispectra

**DOI:** 10.1101/2023.10.26.564193

**Authors:** Franziska Pellegrini, Tien Dung Nguyen, Taliana Herrera, Vadim Nikulin, Guido Nolte, Stefan Haufe

## Abstract

Phase-amplitude coupling (PAC) is a form of cross-frequency coupling in which the amplitude of a fast oscillation is locked to the phase of a slow oscillation. PAC has been proposed as a mechanism for integrating slow large-scale networks with fast-oscillating local processes in the brain. On a signal level, PAC can be observed in single time series, reflecting local dynamics, or between two time series, potentially reflecting a functional interaction between distinct brain sites. To investigate the role of PAC as a mechanism of brain signalling, it is important to distinguish these two cases. However, when mixtures of underlying signals are observed, between-site PAC can spuriously emerge even if the true interaction is only local (within-site). This problem arises in electrophysiological recordings where mixing occurs due to volume conduction or the presence of a shared electrical reference. To address this problem, we propose to estimate PAC using the anti-symmetrized bispectrum (ASB-PAC). It has previously been shown that the cross-bispectrum can be used to measure PAC while efficiently suppressing Gaussian noise, and that the anti-symmetrized bispectra vanish for mixtures of independent sources. However, ASB-PAC has so far not been used to assess the presence of genuine between-site PAC. Using simulations, we here investigate the performance of different algorithms to detect PAC in a mixed signal setting as well as the performance of the same methods to distinguish genuine between-site PAC from within-site PAC. This is done in a minimal two-channel setup as well as in a more complex setting that assesses PAC on reconstructions of simulated EEG sources. We observe that bispectral PAC methods are considerably better at detecting simulated PAC in the volume conduction setting than three conventional PAC estimators. ASB-PAC achieves the highest performance in detecting genuine between-site PAC interactions while detecting the fewest spurious interactions. Using the ASB-PAC could therefore greatly facilitate the interpretation of future PAC studies when discriminating local from remote effects.

We demonstrate the application of ASB-PAC on EEG data from a motor imagery experiment. Additionally, we present an upgraded version of the free ROIconnect plugin for the EEGLAB toolbox, which includes PAC and ASB-PAC metriscs based on bispectra.

## 1. Introduction

Phase-amplitude coupling (PAC) is a form of cross-frequency coupling in which the amplitude of a fast oscillation is correlated with the phase of a slow oscillation (e.g., Canolty et al., 2006; Tort et al., 2010; Hyafil et al., 2015). It has been suggested that PAC facilitates the coordination of neural activity across various spatial and temporal scales by decomposing neural computations into discrete units of activity for cognitive processes like memory, attention, and learning (Canolty and Knight, 2010; Hyafil et al., 2015).These units are aligned with rhythmic external sensory and motor activities (Canolty and Knight, 2010). Further, it has been suggested that functional PAC may be pathologically increased in patients with movement disorders (De Hemptinne et al., 2013; Yin et al., 2022).

Historically, research focused on the detection of PAC within a single signal (e.g., Osipova et al., 2008; Cohen et al., 2009; Voytek et al., 2010; Florin and Baillet, 2015; Watrous et al., 2015; Yin et al., 2022). However, other studies also aimed to investigate the coupling between the phase of a slow oscillation (SO) in one signal and the fast oscillation (FO) of another signal, originating from two spatially different locations (e.g., Voytek et al., 2015; Daitch et al., 2016; Nandi et al., 2019). Tort et al. (2008) report a coupling between the phase of striatal theta and the amplitude of high-frequency oscillations in the hippocampus, and vice versa, in local field potential (LFP) recordings. Maris et al. (2011) showed the presence of PAC between different intracranial electroencephalography (EEG) electrodes. Others have reported functional PAC between different channels of scalp EEG (Schack et al., 2002; Isler et al., 2008; Jirsa and Müller, 2013). In addition, an EEG study (Gong et al., 2021) reported that only PAC between different ICA components was clinically relevant compared to PAC from the same ICA component. These studies suggest that between-site PAC could serve as a mechanism for the integration of spatially distributed neuronal activity (Jirsa and Müller, 2013; Hyafil et al., 2015). Further, it has been suggested that between-site PAC could serve as a gating mechanism that enables spatially distributed networks to coordinate and operate in parallel (van der Meij et al., 2012).

There have been multiple suggestions on how to measure PAC (e.g., Canolty et al., 2006; Penny et al., 2008; Tort et al., 2010; Özkurt and Schnitzler, 2011; Canolty and Knight, 2010; Kovach et al., 2018; Zandvoort and Nolte, 2021). Traditional metrics are based on a process that first extracts the slow and fast oscillation by bandpass-filtering the original signal in the respective low and high frequency bands. Then, the amplitude of the fast oscillation is extracted, e.g., by calculating its envelope with the Hilbert transform. The relation between the phase of the slow oscillation and the envelope of the fast oscillation is then assessed by calculating either the coherence (Colgin et al., 2009), phase-locking value (Lachaux et al., 1999), or correlation (Bruns and Eckhorn, 2004) between the slow oscillation and the amplitude of the fast oscillation. Alternatively, dependencies between phase and amplitudes can be assessed non-parametrically by either calculating the mean vector length modulation index (MI Canolty et al., 2006), or by comparing their joint distribution to a uniform distribution using the Kullback-Leibler Modulation Index (Tort et al., 2010). A downside of all of these metrics is that they heavily depend on filtering parameters in the preprocessing step (Aru et al., 2015; Kovach et al., 2018; Zandvoort and Nolte, 2021). More recent studies have pointed out that the cross-bispectrum or cross-bicoherence can also be employed to assess PAC (e.g., Hyafil, 2015; Kovach et al., 2018; Zandvoort and Nolte, 2021). Indeed, Hyafil (2015) clarified that bicoherence represents phase–phase coupling between three phases and thus essentially corresponds to PAC. Zandvoort and Nolte (2021) further showed theoretically that bispectra closely correspond to conventional PAC metrics for specific filter settings. These studies highlight that metrics based on the bispectrum have two advantages: First, unlike most other PAC metrics, they do not depend of filter settings. And second, the cross-bispectrum efficiently suppresses Gaussian noise (e.g., Nikias and Pan, 1988).

Electrophysiological recordings like intracranial or scalp electroencephalography (EEG), as well as Magnetoencephalography (MEG), are well suited to study PAC because of their high temporal resolution. However, EEG/MEG recordings are often corrupted by noise and signal mixing (Nolte et al., 2004; Haufe et al., 2013; Bastos and Schoffelen, 2016; Schaworonkow and Nikulin, 2021). In case of sensor-level MEG and scalp EEG, mixing arises due to volume conduction, i.e., the superposition of underlying brain sources on the sensors (e.g., Schaworonkow and Nikulin, 2021). To reconstruct the underlying neuronal sources, it is common practice to use inverse models that project sensor-level activity to source locations in the brain (e.g., Baillet et al., 2001). However, since inverse algorithms are in general unable to reconstruct sources perfectly, signal mixing is not entirely eliminated. This phenomenon is called source leakage (Schoffelen and Gross, 2009). Source mixing can also occur in intracranial EEG recordings, like local field potentials (LFP), e.g., due to volume conduction and a shared electrical reference being used for different recording sites (Bastos and Schoffelen, 2016).

Signal mixing can degrade the statistical power for detecting PAC within a single channel of measured or reconstructed activity but it cannot lead to spurious PAC. However, when the goal is to study genuine between-site PAC, i.e., coupling between the phase of a slow oscillation coming from one brain location, and the amplitude of a fast oscillation coming from a different brain location, signal mixing can be a confounding factor. In this case, one may erroneously interpret within-site PAC as between-site PAC when slow and fast oscillation signal components leak into both studied channels (Figure 2). This raises the question: how can we distinguish genuine between-site PAC from spurious between-site PAC arising from signal leakage of sources exhibiting within-site PAC? Note that we use the term ‘between-site’ here to refer to the presence of two anatomically distinct neural current sources projecting to M/EEG sensors with different topographies. This would entail cases where sources are physically close to another but currents have distinct spatial orientations either due to a sharp folding of the cortical mantle in the vicinity of the sources or due to recruitment of different neural subpopulations.

There have been various suggestions on how to solve the problem of signal leakage in the context of functional connectivity (FC). For example, Colclough et al. (2015) proposed a symmetric orthogonalization of the time-domain signals before estimating FC metrics. Shahbazi et al. (2010) suggested to compare FC metrics against a null distribution, generated from permuted independent components (IC surrogate approach). And Chella et al. (2014) suggested to use an anti-symmetrization of the cross-bispectrum to eliminate the effects of mixing artifacts on the bispectrum and illustrated that the anti-symmetrized bispectrum (ASB) vanishes for mixtures of independent sources. To date, however, anti-symmetrized bispectra have not been used to define metrics for genuine between-site PAC (ASB-PAC).

Previous research has established that PAC can be quantified using traditional measures like the MI or through third-order cumulants, such as the bispectrum. However, it remains unclear if these methods can also be used to identify the presence of genuine between-site PAC. Moreover, these metrics are vulnerable to signal mixing (e.g., Chella et al., 2014). In this study, we test three strategies aimed at enhancing the robustness of PAC metrics to signal mixing. These strategies include ASB-PAC, MI based PAC calculated on orthogonalized time series, and the IC surrogate approach. We address the following main questions:

- Are the bispectrum and the MI suitable methods to identify genuine between-site as opposed to within-site PAC?
- Is ASB-PAC a valid measure of between-site PAC?
- Can orthogonalization or the IC surrogate approach be used to make MI basec PAC robust against signal mixing?

In addition to these questions, we are interested in the impact of the signal-to-noise ratio and the number of underlying ground-truth interactions on the accuracy of detection. By answering these questions, we aim at finding an analysis approach that ensures a reliable detection of between-site PAC, even in the presence of mixed noise.

To this end, we first quantify the performance of different algorithms with respect to their ability to detect PAC in a mixed signal setting. Second, we investigate the performance of these methods to distinguish genuine between-site PAC from within-site PAC in the mixed signal setting. We first conduct a set of simple experiments involving only two channels in different signal- and noise settings. These experiments resemble a recording setup of two, potentially invasive, single electrodes. Next, we translate the simulation to an advanced whole-brain setup, reflecting a scalp EEG recording. Finally, we investigate the presence of robust between-site PAC during motor imagery using real scalp-EEG data (Sannelli et al., 2019).

## 2. Methods

### 2.1 Univariate PAC

PAC refers to the coupling between a slow and a fast oscillation. For a given univariate time series *x*(*t*) ∈ ℝ, *t* = 1, …, *N*_*t*_, with *N*_*t*_ denoting the number of time points, the goal is to determine whether there is PAC between the two frequency bands *l* and *h*. If 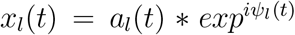 and 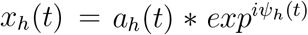 denote the low and high-frequency parts of the univariate signal, with *a*(*t*) ∈ ℝ denoting the amplitude and *ψ*(*t*) ∈ ℝ denoting the phase of the complex-valued signal, PAC refers to the relation between the phase of in the low frequency band *ψ*_*l*_(*t*) and the amplitude envelope of the high frequency band *a*_*h*_(*t*).

### 2.2 Within-site and between-site PAC

PAC can occur either between two oscillations within the same signal *x*(*t*) (*within-site* PAC), or between two oscillations coming from two different signals or sites *x*(*t*) and *y*(*t*) ∈ ℝ (*between-site* PAC). In the latter case, PAC is defined as a relation between the phase of signal *x*_*l*_(*t*) in the low frequency band *ψ*_*lx*_(*t*) and the amplitude envelope of signal *y*_*h*_(*t*) in the high frequency band *a*_*hy*_(*t*).

When measuring PAC between sites in a setting with source mixing, we need to distinguish between three scenarios: Scenario I): There is no PAC between *ψ*_*lx*_(*t*) and *a*_*hy*_(*t*). Scenario II): There is PAC between *ψ*_*lx*_(*t*) and *a*_*hy*_(*t*) that arises from signal mixing, e.g., *x*(*t*) = *c* ∗ *s*(*t*) + *n*_1_(*t*) and *y*(*t*) = *b* ∗ *s*(*t*) + *n*_2_(*t*), with *c* and *b* ∈ ℝ representing two scaling factors, *s*(*t*) ∈ ℝ denoting a time series containing univariate PAC, and *n*_1_(*t*) and *n*_2_(*t*) ∈ ℝ representing two noise time series. Scenario III): There is genuine PAC between *x*(*t*) and *y*(*t*), i.e., PAC that cannot be explained by signal mixing: *x*(*t*) = *c* ∗ *s*_*low*_(*t*) + *n*_1_(*t*) and *y*(*t*) = *b* ∗ *s*_*high*_(*t*) + *n*_2_(*t*), with *s*_*low*_(*t*) and *s*_*high*_(*t*) ∈ ℝ representing two signals that are phase–amplitude coupled.

### 2.3 Methods that estimate PAC

#### 2.3.1 Conventional metrics

##### Modulation Index

The idea behind conventional PAC metrics (Canolty et al., 2006; Tort et al., 2010; Özkurt and Schnitzler, 2011) is to assess the statistical dependence between a phase and an amplitude time series. First, they estimate the distributions of the phase of a time series *x*_s_(*t*) and the amplitude of a second time series *x*_f_ (*t*), which are hypothesized to contain PA-coupled slow and fast oscillations, respectively. In practice, it might not be known a-priori at what frequencies a given pair of time series exhibits PAC. Therefore, exploratory analyses might sweep through a range of sensible frequency combinations, thereby also swapping the roles of the two time series as slow or fast oscillations. The slow and fast signal components *x*_s_(*t*) and *x*_f_ (*t*) are extracted by filtering *x*(*t*) in the low and high frequency bands, respectively. In the next step, the time-dependent phase of the slow signal *ψ*_s_(*t*) is obtained through a Hilbert transform of 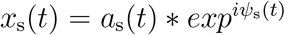. Likewise, the amplitude (envelope) time course of the fast signal, *a*_f_ (*t*), is obtained through a Hilbert transform of 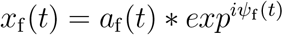.

To obtain the mean vector length MI (Canolty et al., 2006), the two time series are combined into a composite complex-valued signal:

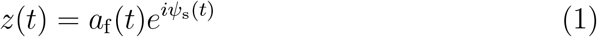

Finally, the MI is obtained as the mean of the absolute value of *z* across time.

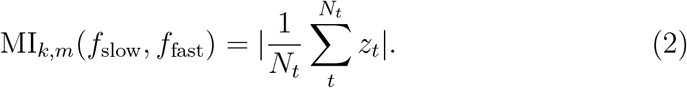

Note that other versions of the MI have been introduced (Tort et al., 2010; Özkurt and Schnitzler, 2011). In brief, the metric proposed by Tort et al. (2010) assesses the degree of the coupling by calculating the Kullback-Leibler distance of the observed phase-amplitude distribution from a uniform distribution. The metric proposed by Özkurt and Schnitzler (2011) extends the original MI by a normalization with the power of the amplitude vector.

Variants of the MI have been used to investigate both within- and between-site PAC; however, it is unclear whether they can distinguish genuine between-site PAC from PAC arising from mixtures of signals exhibiting within-site PAC.

##### Bispectral PAC estimation

The cross-bispectrum belongs to the class of higher-order poly-spectra. Precisely, it is the two-dimensional third-order statistical moment in frequency domain. To estimate the bispectrum, the time series are cut into epochs *e* ∈ 1, …, *N*_*e*_, and Fourier-transformed. In its most general form, it is calculated for a combination of three channels 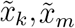, and 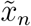, and between two frequencies *f*_1_ and *f*_2_:

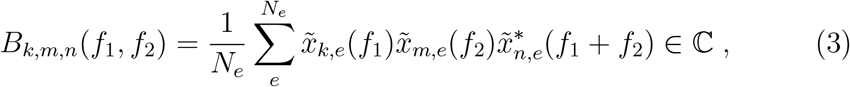

where .^∗^ denotes the complex conjugation.

Recent works have pointed out that PAC can also be estimated from the bispectrum (Kovach et al., 2018; Zandvoort and Nolte, 2021). To this end, the bispectrum *B* is calculated between channel *k* at frequency *f*_1_ and channel *m* at frequencies *f*_2_ and *f*_1_ +*f*_2_, where 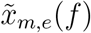 denotes the *e*-th epoch of the Fourier-transformed data of channel *m* at frequency *f* :

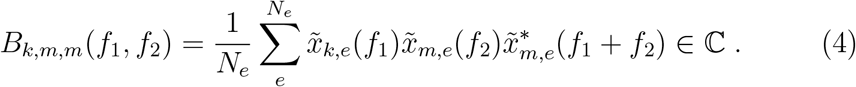

While some works (Kovach et al., 2018; Zandvoort and Nolte, 2021) have shown analytically that the bispectrum is a suitable measure for PAC, it has not been extensively and systematically tested. Moreover, it has not been tested yet whether bispectra can be used to assess genuine between-site PAC not explained by within-site PAC.

##### Filter settings

As discussed by Zandvoort and Nolte (2021), PAC between a slow and a fast oscillation with respective peak frequencies *f*_slow_ and *f*_fast_ corresponds to bispectral interactions at frequency triples 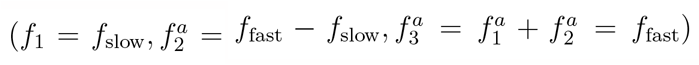 and 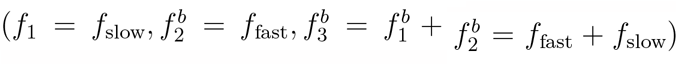 (see Figure 2). Note that the relevance of the “sidelobe” frequencies 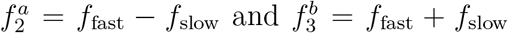 in PAC arises from the multiplicative modulation of the fast signal by the amplitude of the slow signal, which amounts to a convolution in frequency domain. As a result, 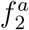 and 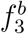 are visible also as additional peaks of spectral power (sidelobes) around the peak of the fast oscillation.

For conventional metrics such as the MI, there have been discussions about how to correctly set the filter parameters, and especially the band width to obtain the fast oscillation (e.g., Berman et al., 2012; Aru et al., 2015; Zandvoort and Nolte, 2021). For example, some works have recommended to set the filter broad enough to include both side lobes 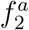 and 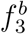. However, Zandvoort and Nolte (2021) pointed out that this leads to a smeared estimation of PAC and instead recommend to include only *f*_fast_ and one of the two side lobes. Here we propose a third alternative—analogous to the bispectral estimate—namely to estimate PAC twice, once with filter settings that include 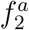 and *f*_fast_, and with filter settings that include *f*_fast_ and 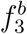.

Note, however, that the notations typically used for bispectral and conventional PAC metrics are not consistent (Zandvoort and Nolte, 2021): while the bispectrum is written as a function of two frequencies *f*_1_ and *f*_2_, implying that the third frequency is fixed at *f*_1_ + *f*_2_, the MI is typically written as a function of the slow frequency and the *center* frequency of the fast oscillation, assuming a symmetrical filter around it. Thus, bispectral PAC at [*f*_slow_, *f*_fast_] corresponds to MI-PAC at [*f*_slow_, *f*_fast_ + 0.5*f*_slow_].

Based on these considerations, and following the conventional notation, the final bispectral PAC estimates in this study are obtained as

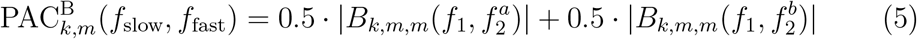

and the final MI-PAC estimates are obtained as

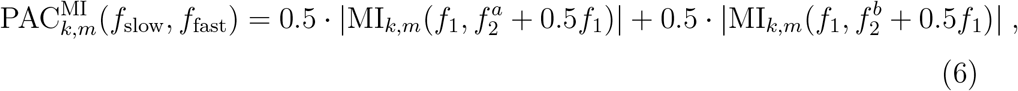

where the band width of the fast-frequency filter is not much larger than *f*_fast_ − *f*_slow_ (see Section 3).

### 2.4 Statistical testing

To statistically assess whether an observed MI is statistically significant, the true MI can be compared against an empirical null distribution. Samples consistent with the null hypothesis of no PAC being present can be obtained by randomly permuting the order of the epochs of one of the time series and subsequently recalculating PAC on the permuted epochs:

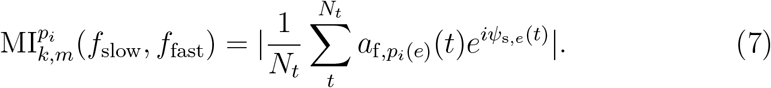

for the *i*th iteration with *i* = 1, …, *S*, where *p*_*i*_(*e*) is a random permutation of the epochs *e* = 1, …, *N*_*e*_.

Analogously, for the bispectral method, the epochs of the Fourier-domain data 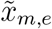 can be permuted to obtain a null distribution for statistical testing:

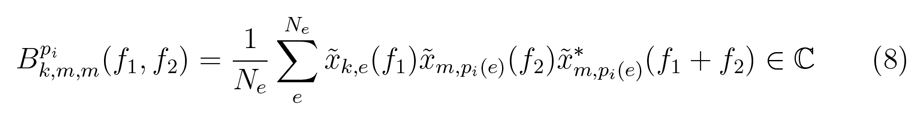

This procedure is repeated *S* times to obtain the desired surrogate distribution. P-values are then calculated as follows:

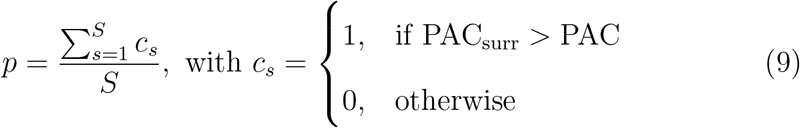

### 2.5 Conventional metrics and between-site PAC

Neither the bispectrum nor the MI are not able to distinguish genuine between-site PAC from spurious PAC arising from signal mixing (Scenario II vs. Scenario III described in Section 2.2). That is, neither metric is able to identify whether observed PAC originates from a single underlying source that has leaked into the two time series, or from two different underlying sources. Similarly, the permutation-based statistical assessment that we propose to test whether the MI or bispectrum is significant is unable to differentiate between within-site PAC and between-site PAC. This is because permutations disrupt all data interactions, including those caused by mixed sources. As a consequence also interactions from the type of Scenario II are tested significant. To address this limitation, alternative methods are essential. In the following, we summarize three approaches that are candidates to solve this problem.

### 2.6 Anti-symmetrization

For the bispectrum, anti-symmetrization has been proposed to correct for effects that arise from signal mixing (Chella et al., 2014). Based on anti-symmetrized bispectra, we can define the following PAC metric whose expected value for mixtures of independent (potentially within-site PAC coupled) signals is zero:

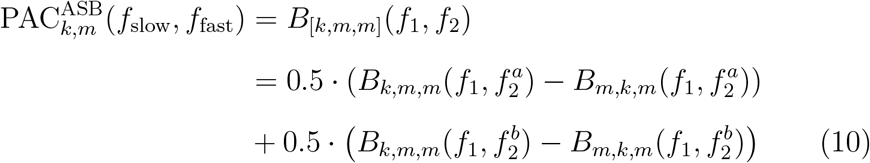

If the slow and fast oscillations originate from two distinct sites, implying a genuine between-site interaction, then we would expect to observe the slow oscillation solely at one site and the fast oscillation at the other site. Contrarily, if the PAC interaction is derived from a single source that leaks to two sites, then both the slow and fast oscillations should be present at each site. Taking advantage of this fact, we can discern genuine interactions based on their anti-symmetric nature by subtracting the symmetric part of the interaction. However, it is unclear whether this leakage-corrected bispectrum can still be interpreted as PAC. Therefore a systematic evaluation whether the ASB-PAC can be used to estimate between-site PAC is needed.

### 2.7 Orthogonalization

An equivalent technique to anti-symmetrization for bispectra is not available for the MI. However, several methods have been proposed in the literature as general solutions to remove the effect of source mixing from data or from statistical contrasts against null distributions. First, a multivariate symmetric orthogonalization technique has been proposed by Colclough et al. (2015). This technique removes instantaneous correlations between multivariate time series, which is proposed as a general correction for source leakage effects in functional connectivity analyses. The authors demonstrate its usability in a simulation where partial correlations between power envelopes are analyzed, as well as in a real resting-state MEG data example. Thus, even though symmetric orthogonalization is not specifically designed for PAC problems, it may be suitable to solve the source mixing problem also for between-site PAC.

The advantage of the symmetric orthogonalization over similar decorrelation schemes (e.g., Hipp et al., 2012; Brookes et al., 2012) is that the result is not dependent on the ordering of the region time series, and that the resulting time series are as close as possible (in the least-squares sense) to the original time series. This orthogonalization scheme is based on the Löwdin method (Löwdin, 1950) and can be implemented using a singular value decomposition (Colclough et al., 2015; Annavarapu, 2013):

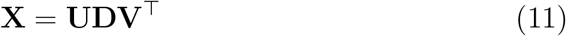

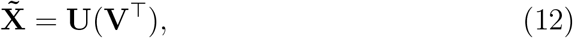

where 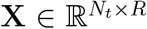 contains *R* time series to be orthogonalized. *R* indicates the number of time series that are investigated, and **V**^⊤^ denotes the transpose of **V**. Using the ‘economy version’ of the SVD, 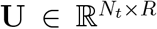 is a matrix of orthonormal time courses, **V** ∈ ℝ^*R*×*R*^ is a matrix of singular vectors, **D** ∈ ℝ^*R*×*R*^ is a matrix of singular values, and 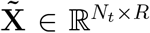 is the matrix of symmetrically orthogonalized time courses.

To test the statistical significance of the resulting PAC, surrogate data without PAC are constructed. To this end, the epochs of one times series are permuted and concatenated back to form a time series. Then the symmetric orthogonalization is applied to the concatenated time series, and PAC 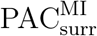 is estimated on the surrogate data. Finally the p-value is calculated as described above.

### 2.8 Surrogate data using independent component analysis

Another general approach to account for artifacts of source mixing is to generate a null distribution by permuting the epochs of independent component (IC) time series (Shahbazi et al., 2010). In this study, this method will be called IC surrogate analysis (IC-surr). The idea of this approach is to construct surrogate data that are statistically as close as possible to the original data, but are still physically realistic mixtures of independent sources rather than being completely independent. The comparison of the observed FC to the distribution of FC obtained from surrogate data can then reveal whether observed FC is a genuine effect that cannot explained by source mixing alone. The authors demonstrate the usablility for the linear measures coherence and imaginary part of coherency, 1:2 phase coupling, and Granger causality in simulated and real EEG data. They show that results obtained with the robust FC measure imaginary coherency cannot be explained by the surrogate data. Further, for Granger Causality, a non-robust measure, they observe that the ground-truth interactions that they simulated were attenuated but not removed in the surrogate data. The authors propose that the IC-surr approach is applicable to other interaction measures as well. Therefore, we also include this approach in our study and test whether it can be used to target the between-site PAC estimation problem.

We here describe the approach exemplarily for a setting with *N*_*s*_ channels. The first step in the IC surrogate analysis is to perform independent component analysis (ICA) of the time series matrix 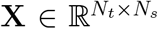. In brief, ICA finds a weighting matrix 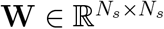 that unmixes the sensor data into ICs that are minimally statistically dependent according to some criterion:

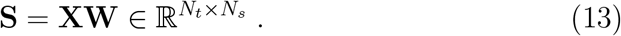

In the present study, we use the Infomax ICA algorithm (Bell and Sejnowski, 1995) as implemented by the `runica.m` function of the EEGlab package (Delorme and Makeig, 2004), but other algorithms are conceivable. Sub-sequently, the ICs are permuted by temporally shifting them randomly with respect to each other. Finally, the concatenated component time series are projected back to the original space:

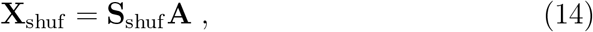

with 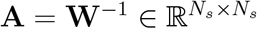 denoting the ICA mixing matrix. From **X**_shuf_, the MI can be calculated again. The p-value is then calculated by comparing the original MI against the IC-surrogate MI distribution.

## 3. Experiments

To test whether the various PAC metrics and robustification approaches are able to distinguish genuine between-site PAC while rejecting contributions from within-site PAC, we performed a number of experiments, which are described below.

First, to ensure that performance differences between the MI- and bispectrum-based PAC metrics cannot be attributed to different performances in the detection of univariate PAC, we conducted a small baseline experiment. To this end, we simulated univariate PAC within a single channel, added noise with varying signal-to-noise ratios (SNRs), and measured PAC between the phase of the slow oscillation and the amplitude of the fast oscillation within the single channel.

We then conducted two experiments involving only *N*_*s*_ = 2 channels, which allow us to quantify the performance of different analysis pipelines in different SNR settings. In 2CHAN-BI (c.f. Table 1), we simulated true bivariate PAC between the two channels. These experiments allow us to study the sensitivity of our analysis pipelines with respect to detecting ground-truth between-channel interactions as a function of SNR. In the 2CHAN-UNI experiments, we simulated two univariate phase–amplitude coupled signals mixed into each of the two channels. This setup enables us to assess false-positive rates of the different approaches. Please note that we here use the term *bivariate PAC* to describe ground-truth between-region or between-channel PAC, reflecting a true interaction between a slow oscillation in one brain source and a fast oscillation in another brain source. In contrast, we use the term *univariate PAC* describing ground-truth within-region or within-channel PAC, reflecting PAC between a slow and a fast oscillation within the same source site.

**Table 1:** Overview of the univariate baseline experiment (1CHAN-UNI) and the differences between the four 2CHAN experimental settings: modeled interaction and performance evaluation (TPR: true positive rate, TNR: true negative rate).

| Simulation |  | Interaction | Evaluation |
| --- | --- | --- | --- |
| 1CHAN-UNI | Single-channel<br>baseline | Univariate PAC within<br>a single channel. | TPR |
| 2CHAN-BI-IND | Two-channel<br>bivariate,<br>independent | Bivariate PAC between<br>two channels. | TPR |
| 2CHAN-BI-MIX | Mixed |  |  |
| 2CHAN-UNI-IND | Two-channel<br>univariate,<br>independent | Two channels with two<br>univariate PAC time<br>series. | TNR |
| 2CHAN-UNI-MIX | Mixed |  |  |

**Table 2:** Overview of the differences between the six whole-brain EEG experimental settings: modeled interaction and performance evaluation (SNR: signal-to-noise ratio, *N*_*I*−*bi*_ number of bivariate interactions, *N*_*I*−*uni*_ number of univariate interactions, PR: percentile rank, gFPR: global false positive rate).

| Simulation |  | Interaction | Evaluation |
| --- | --- | --- | --- |
| EEG-BI | Whole-brain EEG<br>bivariate | Three bivariate interactions between six regions.<br>Other regions contain independent brain noise. | PR |
| EEG-BI-SNR | Varying SNR | Three bivariate interactions between six regions. |  |
| EEG-BI-INT | Varying $N_{I-bi}$ | One, three, or five bivariate interactions. | |
| EEG-UNI | Whole-brain EEG<br>univariate | One univariate PAC time series.<br>Other regions contain independent brain noise. | gFPR |
| EEG-UNI-SNR | Varying SNR | One univariate PAC time series. |  |
| EEG-UNI-INT | Varying $N_{I-uni}$ | One, three, or five univariate PAC time series. | |

Subsequently, we considered a whole-brain scalp-EEG setting. From this setting, we expect to gain insights beyond the minimal two-channel setup into how well the methods might work on EEG data in practice. Again, we simulated two cases. In Experiment EEG-BI (c.f. 2), three true bivariate PAC interactions between six randomly selected regions of the brain were modeled, while all other regions emitted only independent brain noise activity. This experiment allows us to study the sensitivity of the tested approaches in a setting with realistic signal mixing. In Experiment EEG-UNI, we considered three active regions containing independent univariate within-region PAC, while again all other regions elicit independent brain noise only. This experiment is important to test the suitability of the approaches in eliminating spurious PAC detection in a setting with realistic signal mixing. In the following, we first describe two-channel experiments before moving on to more complex whole-brain EEG experiments. Matlab code to reproduce all experiments is provided^1^. The implementation for the bispectra is based on the open-source METH toolbox^2^.

### 3.1. Baseline experiment on univariate PAC detection

#### 3.1.1. Data generation

We generated time series at a sampling rate of 200 Hz with a recording length of *N*_*t*_ = 120 000 samples amounting to a duration of 10 minutes. The signal was generated as random white Gaussian noise filtered in the low frequency band (defined here as ranging from 9 to 11 Hz) for the slow oscillation, denoted *x*_s_(*t*) with *t* ∈ {1, …, *N*_*t*_}, and in the high frequency band (ranging from 58 to 62 Hz) for the fast oscillation, *x*_f-raw_(*t*). Throughout, we used zero-phase forward and reverse second-order digital Butterworth band-pass filters. The PAC interaction was modeled as coupling between the phase of the slow oscillation and the amplitude of the fast oscillation. To achieve this, we extracted the phase of the slow oscillation *ψ*_s_(*t*) and the phase of the fast oscillation *ψ*_f_ (*t*) by means of the Hilbert transform, from which we calculated the modulated fast oscillating signal *x*_f_ :

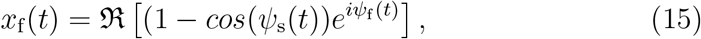

with ℛ [*x*] denoting the real part of *x*. We divided both the slow and the fast oscillating signal by their -norms for normalization: 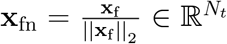 and 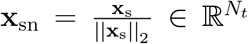, where 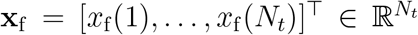 and 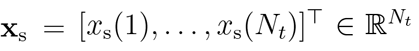 are the concatenated values of *x*_f_ (*t*) and *x*_s_(*t*) at all time points, respectively. Finally, we summed up the slow and the fast oscillation to yield the univariate PAC signal:

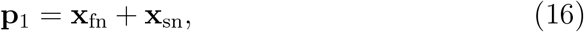

and then divided the signal by its Frobenius norm: 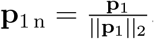.

We generated a time series 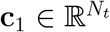, representing channel noise, as random white noise. The channel noise time series was then also divided by its _2_-norm: 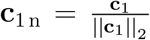. Finally, we formed a weighted sum of signal and channel noise:

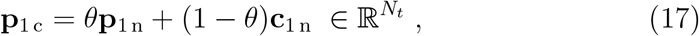

where the parameter *θ* ∈ [0, 1] defines the signal-to-noise ratio on channel level (SNR_c_), which can be expressed in decibel (dB) as: 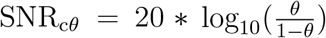. We compared the performance of various MI variants and the bispectrum without anti-symmetrization in detecting the univariate PAC for the following SNR_c_ values: noise only (*θ* = 0), -19 dB (*θ* = 0.1), -12 dB (*θ* = 0.2), -7 dB (*θ* = 0.3), and -4 dB (*θ* = 0.4).

#### 3.1.2 PAC analysis

To estimate PAC, we first cut the time series into epochs. PAC estimation based on the MI requires one to cut both signals into long epochs (Tort et al., 2010). That is, for MI-based metrics, we split the data into *N*_*e*_ = 60 epochs of length *L*_*e*_ = 2000 = 10 sec. For bispectral PAC metrics, we cut the time series into *N*_*e*_ = 300 epochs of 2 seconds length (i.e., 400 samples).

To estimate the MI, we filtered the epoched time series in the low and high frequency bands. Since we modelled the ground-truth interaction at *f*_slow_ = 10 Hz and *f*_fast_ = 60 Hz, we calculated the MI once between the slow signal *x*_s_(*t*) filtered between 9 and 11 Hz and the fast signal *x*_f_ (*t*) filtered between 49 and 61 Hz, and once between the slow signal and the fast signal filtered between 59 and 71 Hz (c.f. Section 2.3.1 for a detailed explanation on filter settings). Subsequently, the two MI estimates were averaged.

To estimate the bispectral metrics, we Fourier-transformed every epoch after multiplying it with a 400-point symmetric Hanning window. Afterwards, we estimated the bispectra once at (*f*_1_ = 10 Hz, 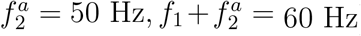) and once at (*f*_1_ = 10 Hz, 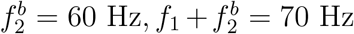). Subsequently, the two estimates were averaged.

For the baseline experiment, we also assessed the performance of additional variants of the MI as introduced by Tort et al. (2010) and Özkurt and Schnitzler (2011). Tort et al. (2010) tested their MI metric against other measures of PAC, like, e.g., the phase locking value, and found that it is suitable to measure univariate PAC. Özkurt and Schnitzler (2011) also compared their proposed MI metric to other PAC measures like the general linear model approach and showed that it is superior to them.

To assess the statistical significance of the PAC estimate, we generated a surrogate distribution with *S* = 1000 samples, from which we calculate a p-value as described in Section 2.4.

#### 3.1.3 Performance evaluation

The experiment was repeated *D* = 100 times to obtain 100 p-values *p*_*d*_. PAC was considered statistically significant for p-values below an *α*-level of 0.05. To assess whether the studied PAC detection pipelines are able to correctly detect the presence of PAC, we evaluated the true positive rate (TPR):

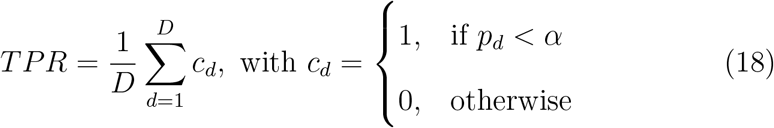

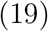

### 3.2 Two-channel experiments

In the two-channel experiments (Figure 1), we generated either two underlying time series exhibiting a true bivariate PAC interaction (referred to as 2CHAN-BI), or two independent time series each exhibiting univariate PAC (referred to as 2CHAN-UNI, see Section 3.2.1). The underlying signals were then further linearly mixed into two measurement channels (see Figure 2 for an overview of the generated signals). The objective of the two-channel setup is to provide a simplified context for showcasing the characteristics of the different PAC estimation approaches. We expect that all non-robust metrics are not capable of distinguishing within-from between-site PAC. Conversely, the ASB-PAC should be able to distinguish these. However, it is unclear how the orthogonalization approach (Colclough et al., 2015) and the IC surrogate method (Shahbazi et al., 2010) might perform. The two-channel setup offers a good way to analyze their behavior in a simple controlled setup.

**Figure 1:**
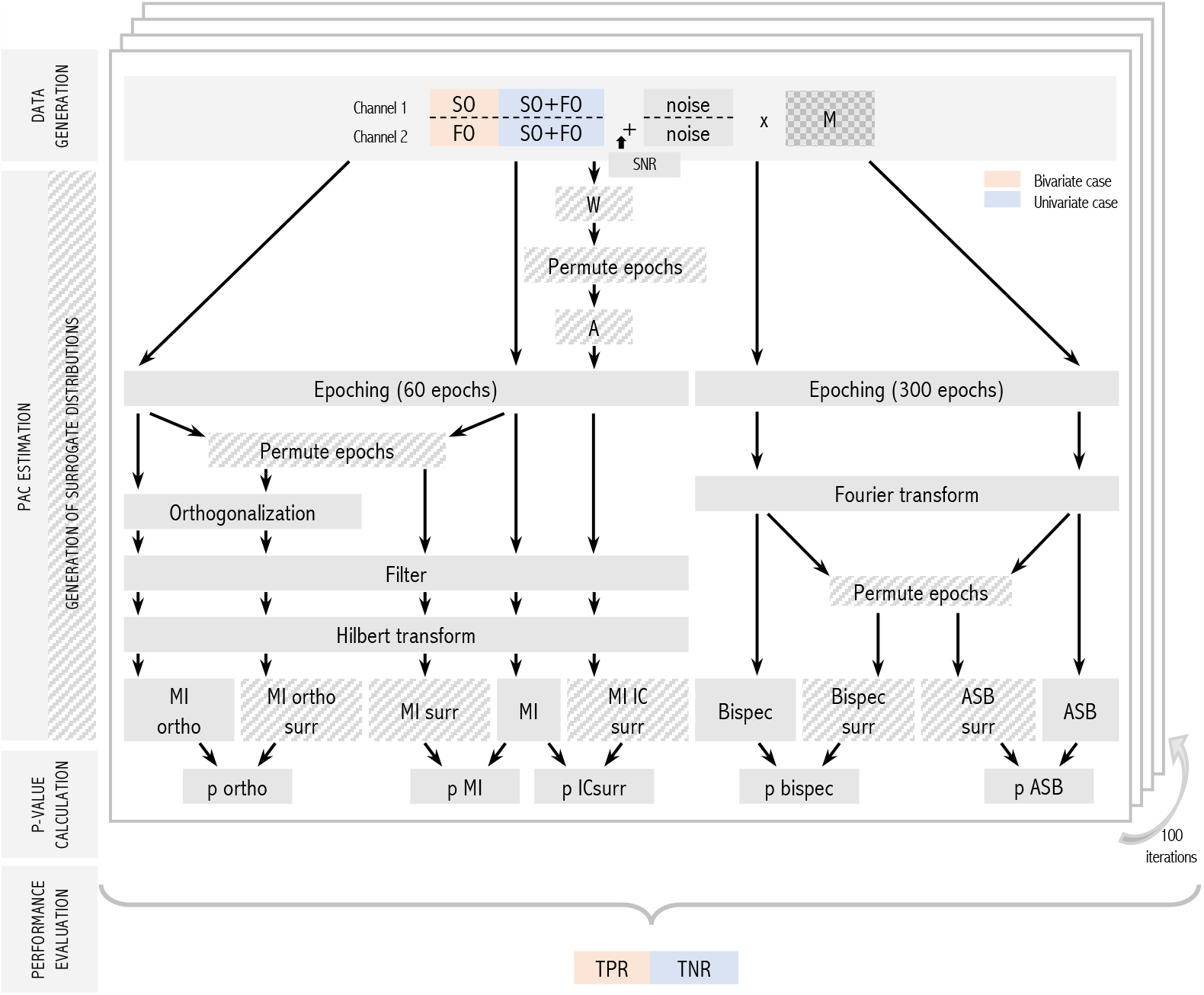
Structure of the two-channel experiments and data analysis consisting of the following steps: data generation, PAC estimation and generation of surrogate distributions, p-value calculation, performance evaluation. SO: slow oscillation, FO: fast oscillation, M: mixing matrix, ICs: components retrieved from an independent component analysis (ICA), W: ICA unmixing matrix, A: ICA mixing matrix, MI: modulation index, ASB-PAC: anti-symmetrized bispectrum, TPR: true positive rate, TNR: true negative rate.

**Figure 2:**
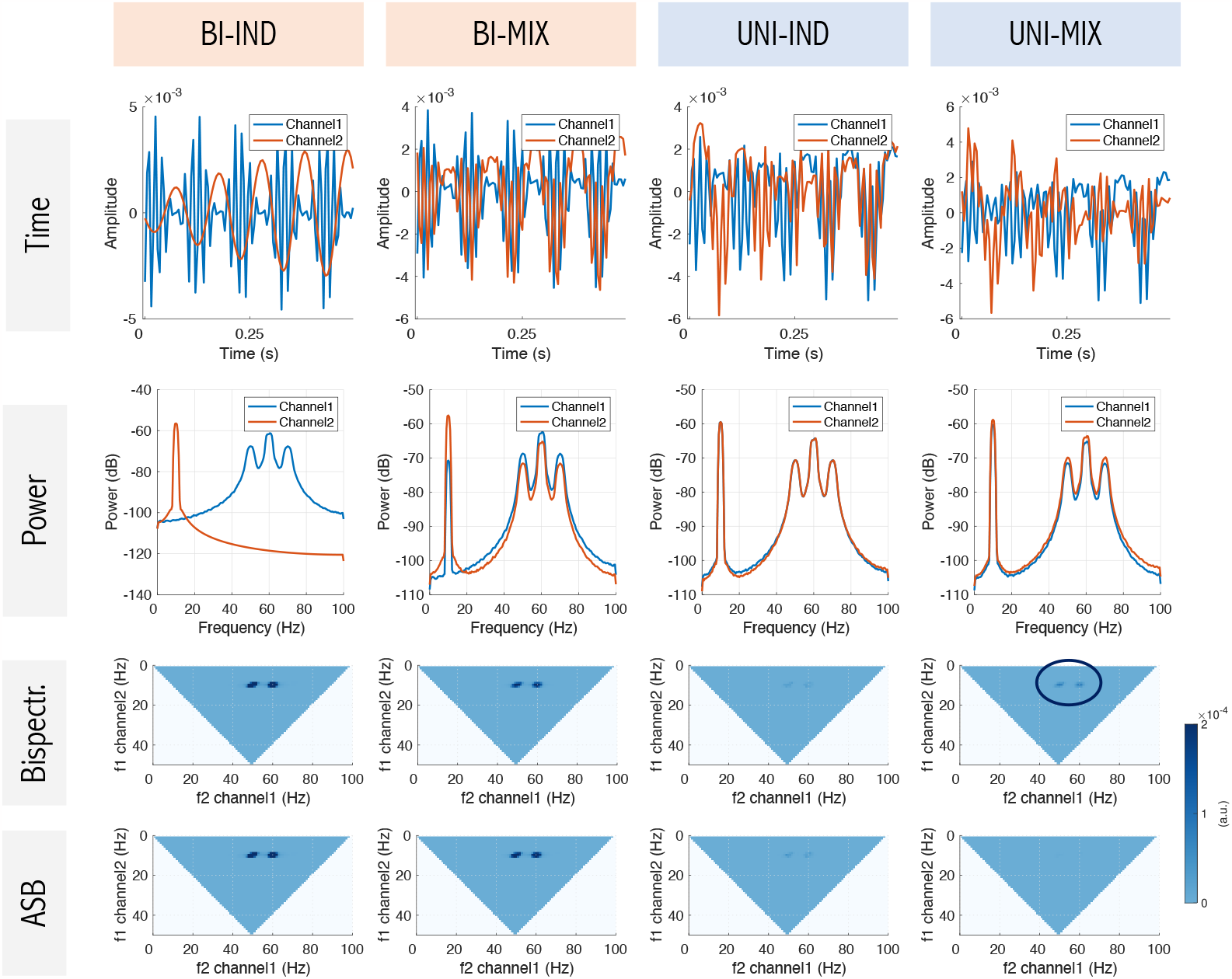
Overview of signals in the 2CHAN experiments. Shown are exemplary signals without the addition of noise. First row: excerpt of signal in time domain. Second row: power spectrum. Third row: full bispectrum. Fourth row: full anti-symmetrized bispectrum. Substantial bispectral energy is observed in the UNI-MIX case despite the absence of between-site PAC interactions (black circle).

#### 3.2.1 Data generation

As in the baseline experiment, we generated a slow oscillation (9 to 11 Hz) and a fast oscillation (58 to 62 Hz) time series at a sampling rate of 200 Hz with a recording length of *N*_*t*_ = 120 000 samples = 10 minutes. As described above, the amplitude of the fast oscillation was modulated by the phase of the slow oscillation. Afterwards, they were normalized by their _2_-norms.

We generated time series **c**_1_ and 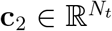, representing noise in the two channels, as random white noise. The channel noise time series were then also divided by their _2_-norms: 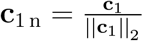 and 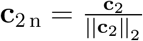.

In the experimental setting comprising an underlying signal with bivariate PAC (2CHAN-BI), we added the channel noise to the slow and the fast oscillating signals, respectively:

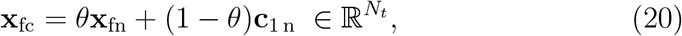

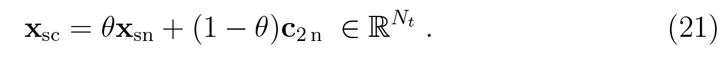

To simulate signal mixing (MIX), we multiplied the two channel time series by a mixing matrix **M**: **d**_bivar_ = **x** ∗ **M**, with 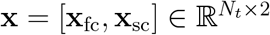, and with

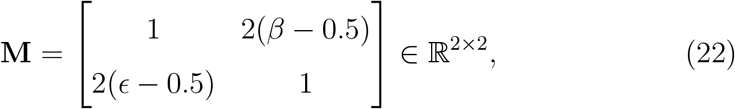

with *β* and *ϵ* representing two random parameters drawn from a uniform distribution on the open interval (0, 1).

In the setting comprising two underlying signals with univariate PAC (2CHAN-UNI), we generated two independent univariate PAC signals **p**_1_ and 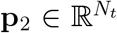, representing the data of the two channels. To generate data of the first channel, we summed up the slow and fast oscillation:

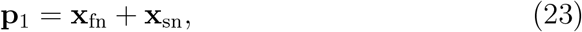

then divided the result by its Frobenius norm: 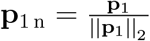, and finally added channel noise:

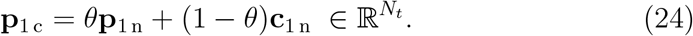

Data for the second channel were generated analogously. Finally, we again mixed the two channels with the mixing matrix **M**: **d**_univar_ = **p** ∗ **M**, with 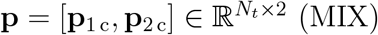 (MIX).

We conducted experiments for the following SNR_c_s: noise only (*θ* = 0), -12 dB (*θ* = 0.2), -4 dB (*θ* = 0.4), 4 dB (*θ* = 0.6), and 12 dB (*θ* = 0.8). To quantify the effect of signal leakage on the different PAC metrics, we conducted all experiments once without mixing (*β* = *ϵ* = 0.5, denoted as 2CHAN-BI-IND, 2CHAN-UNI-IND), and once with the additional random linear mixing of the two source channels (2CHAN-BI-MIX, 2CHAN-UNI-MIX).

The PAC analysis of the two-channel experiments follows the simulation flow described in Section 3.1 (1CHAN-UNI). Note that for both the true PAC score (PAC) and the PAC scores (PAC_surr_) obtained from surrogate data, we estimated the coupling between the two time series in both directions, but always selected the larger PAC score of the two for further processing.

#### 3.2.2 Performance evaluation

To assess whether the studied PAC detection pipelines are able to correctly detect the presence of bivariate (between-site) PAC in experimental setting 2CHAN-BI, we evaluated the TPR. Conversely, in setting 2CHAN-UNI, in which only univariate (within-site) PAC was simulated, we evaluated the true negative rate (TNR) to assess whether metrics correctly reject the hypothesis that the observed PAC originates from two different sources.

Every experiment was repeated for *D* = 100 times to obtain 100 p-values *p*_*d*_. PAC is considered statistically significant for p-values below an *α*-level of 0.05. The TPR (for *θ >* 0 and TNR (for *θ* = 0 were calculated as follows:

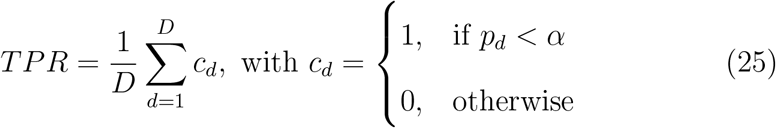

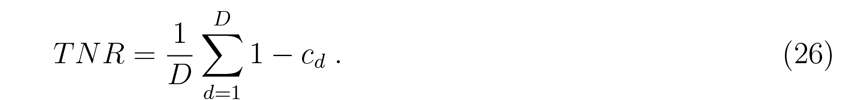

### 3.3 Whole-brain EEG experiments

To assess the extent to which the considered PAC estimation pipelines are also able to distinguish between-from within-site PAC in a practical setting involving source mixing, we next conducted a set of advanced EEG experiments (Figure 3). This included the presence of noise sources, source mixing through a realistic EEG forward model, realistic preprocessing including source reconstruction, and the application of dimensionality reduction techniques. Further, we here varied the SNR and the number of ground-truth interactions to assess their influence on the sensitivity and specificity of the candidate PAC detection pipelines.

**Figure 3:**
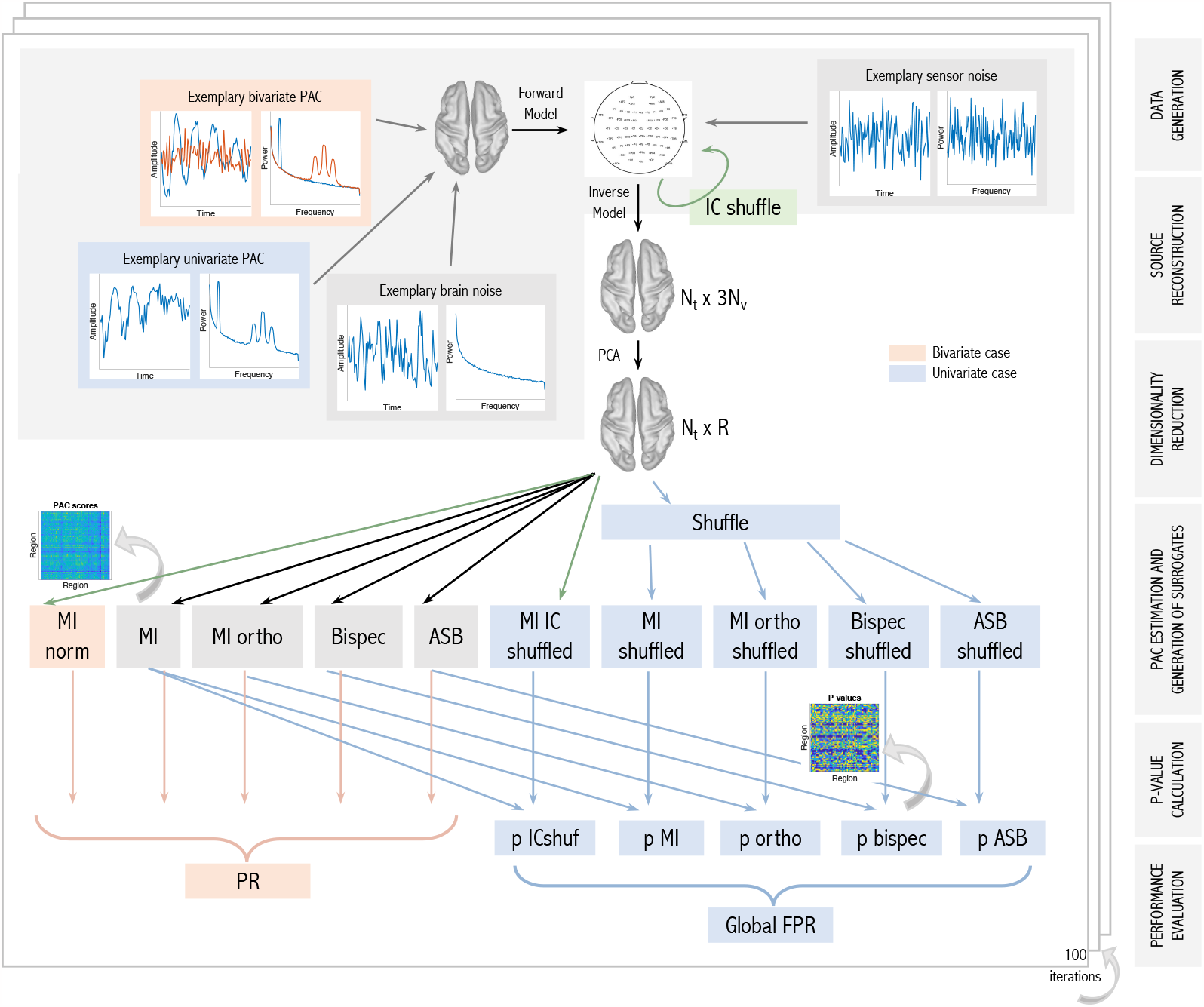
Structure of the whole-brain EEG simulation study consisting of the following steps: data generation, source reconstruction, dimensionality reduction, PAC estimation and generation of surrogates, p-value calculation, and performance evaluation. In case of genuine bivariate (between-site) simulated PAC, detection performance is evaluated with the percentile rank (PR). In the case of solely univariate (within-site) PAC, null distributions are generated and a p-value for every region combination is calculated. From this, we calculate the global false positive rate (global FPR).

#### 3.3.1 Data generation

We generated time series at a sampling rate of 200 Hz with a recording length of *N*_*t*_ = 120 000 samples, amounting to 10 minutes duration. The slow and fast oscillations *x*_s_ and *x*_f_ were generated as in the two-channel experimental setup (see Section 3.2.1). Additionally, to make the signal more realistic, we here transformed *x*_s_ and *x*_f_ to a 1/f-shape before further processing. To this end, the signal was first Fourier-transformed. Subsequently, every value of the Fourier-transformed signal was divided by its corresponding frequency. Afterwards, the signal was transformed back to time domain by using an inverse Fourier transform.

In the *bivariate* PAC Experiments EEG-BI, we generated bivariate interactions between *N*_*I*−*bi*_ pairs of time series, as described in Section 3.2.1. We then added pink (1/f scaled) background noise as follows: For every interaction, we generated time series **b**_1_ and 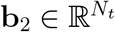, representing the background noise for the two sources. The background noise time series were then also divided by their _2_-norms: 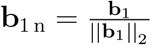 and 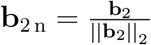.

We added the background noise to the slow and the fast oscillating signals, respectively:

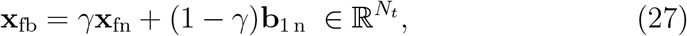

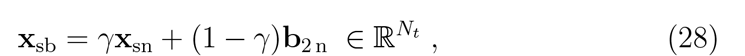

where *γ* ∈ [0, 1] defines the signal-to-noise ratio on source level (SNR_s_) and is fixed here to *γ* = 0.6.

In the *univariate* PAC experiments EEG-UNI, we generated *N*_*I*−*uni*_ univariate PAC time series by summing up the PA-coupled slow and fast oscillations to form a single signal, and by adding pink background noise as described above.

In both the bivariate and univariate PAC experiments, we additionally simulated activity of non-interacting sources—referred to as *brain noise*— using mutually independent random pink noise signals only without additional activity in the alpha band or any other specific frequency band.

#### 3.3.2 EEG signal simulation

EEG forward modeling was carried out in Brainstorm (Tadel et al., 2011) using the ICBM152 anatomical head template, which is a non-linear average of the magnetic resonance (MR) images of 152 healthy subjects (Mazziotta et al., 1995). Within the ICBM152 anatomical model, *N*_*v*_ = 1895 dipolar sources were placed in the cortical gray matter half-way between the white matter—gray matter and gray matter—cerebro-spinal fluid interfaces. Each dipole thereby models the net primary electrical current elicited by a large population of pyramidal cortical neurons. A cortical parcellation according to the Desikan-Killiany atlas (Desikan et al., 2006) was used to assign each dipole to one of *R* = 68 regions. A set of *N*_*s*_ = 97 EEG sensors were placed on the scalp following the standard BrainProducts ActiCap97 channel montage. The mapping from dipolar sources to EEG sensors was calculated with a three-shell boundary element (BEM) model using the OpenMEEG (Gramfort et al., 2010) package with the three shells representing the brain-skull, skull-skin and skin-air interfaces, respectively. The result is summarized in the *leadfield* matrix 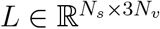.

For the purpose of this simulation, a single ground-truth source time series was placed in each of the 68 regions of the DK atlas, where the location within each region was chosen at random. The spatial orientation of all sources was set to be perpendicular to the cortical surface at the given location. In the univariate PAC experiments, the regions in which the univariate PAC sources are placed were chosen randomly. In the bivariate PAC experiments, the region pairs in which the pairs of interacting PAC signals were placed were also chosen randomly, where we ensured that low and high-frequency components were not located within the same region. In all other regions of the DK atlas, a single brain noise source was placed. The following steps were described extensively in (Pellegrini et al., 2023). In brief, signal and brain noise sources were separately projected to the EEG sensor space using the lead field matrix. On sensor level, channel noise was generated as Gaussian white noise (c.f. Section 3.2.1), and mixed with the brain noise (brain noise to sensor noise ratio of 19 dB) to obtain the *total noise*. Afterwards, we summed up signal and noise with a predefined total SNR, denoted_t_. We adjusted the default total SNR_t_ to 12 dB in the univariate PAC experiments, and to 0 dB in the bivariate PAC experiments. These different settings were chosen to create a challenging problem setting in both cases. As a last step, to make the generated data more realistic, we high-pass filtered the generated sensor data with a cutoff of 1 Hz. The resulting time series on sensor level is here called 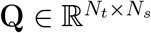.

If not indicated otherwise, all experiments with ground-truth bivariate PAC interactions had the following default settings:

- SNR_t_ = 0 dB
- Number of PAC interactions: *N*_*I*−*bi*_ = 3.

All experiments simulating univariate PAC sources within individual regions had the following default settings:

- SNR_t_ = 12 dB
- Number of PAC seed regions *N*_*I*−*uni*_ = 1.

In Experiment EEG-UNI-SNR, we assessed PAC detection performance as a function of SNR, considering SNRs of 0 dB, 7.4 dB, and 12 dB. The SNRs for Experiment EEG-BI-SNR was chosen to be lower to avoid ceiling effects in the results: here, we assessed the metrics’ performances for SNRs of -7.4 dB, 0 dB, and 7.4 dB. In Experiment EEG-BI-INT and EEG-UNI-INT, we tested the influence of the number of ground-truth interactions and show results for 1, 3, and 5 interactions.

In the EEG-BI experiments, ground-truth interacting regions (two disjoint sets of region indices were drawn randomly from a uniform distribution between 1 and *R* in each iteration. In the EEG-UNI experiments, ground-truth seed region(s) containing the univariate PAC signal (seed region index uniformly drawn between 1 and *R*) were drawn randomly. Furthermore, the ground-truth active voxel(s) within regions (uniformly drawn between 1 and the number of sources within the ground-truth region), brain noise and sensor noise, as well as the signal were generated based on (filtered) random white noise processes as described above.

#### 3.3.3 EEG source reconstruction

The following steps and parameter choices build on the results of Pellegrini et al. (2023). We estimated the activity of the underlying dipolar electrical current sources by constructing linearly-constrained minimum variance (LCMV, Van Veen et al., 1997) beamformers 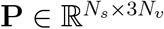 and applying them to the sensor time series:

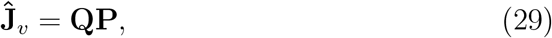

with 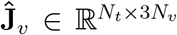 representing the source-level time series. The result is one time series for each of the three spatial orientations of each dipole. To aggregate the reconstructed time series of all dipolar sources within one region, we performed singular value decomposition (SVD), and selected only the strongest SVD component for every region for further processing (c.f. Pellegrini et al., 2023). Let 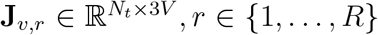, *r* ∈ {1, …, *R*} denote the source time courses of *V* sources within the *r*-th single region. These time courses were aggregated into a single time series by projecting **J**_*v,r*_ onto its strongest SVD component using the filter 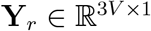:

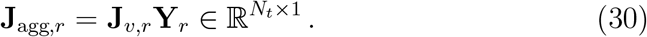

Note that beamformers and PCA that were calculated on unfiltered data comprising both slow and fast signal components in each experimental run.

All region time courses **J**_agg,*r*_, *r* ∈ {1, …, *R*} were then concatenated into a matrix 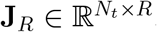, whose columns contain the single time courses **J**_agg,*r*_ of all *R* regions.

#### 3.3.4 PAC analysis

We included the same PAC metrics as in the two-channel experiments in the comparison. In the following, we describe how PAC metrics are applied in the bivariate and univariate whole-brain EEG experiments.

We calculated PAC scores between all region combinations. Note that for the orthogonalization method, we symmetrically orthogonalized all 68 region time series to each other before calculating the MI. Thus, **X** in Eq.(11) now refers to the region time series **J**_*R*_ and *R* = 68 denotes the number of regions.

Here we focus on the ability of the PAC estimation approaches to discriminate between-from within-site PAC in the current whole-brain *bivariate* PAC experiments. Therefore, we did not perform statistical testing for significant PAC like in the two-channel experimental setting. That is, instead of comparing PAC scores to null distributions obtained on permuted data, we directly evaluated the detection performance of PAC metrics with the percentile rank (PR; see Section 3.3.5). An exception was the surrogate data method of Shahbazi et al. (2010), which relies on statistical testing against a suitable null hypothesis, and whose usage is described in the next paragraph. Only in the EEG-UNI experiments, we did generate null distributions by permuting epochs, analogous to what is described for the two-channel experiments. We use the null distributions to calculate a p-value to assess the false positive rate of each approach (see Section 3.3.5).

##### Application of ICA-based surrogate data in whole-brain experiments

Rather than defining its own PAC metrics, the surrogate data approach of Shahbazi et al. (2010) is based on a comparison of observed FC scores against a distribution of FC scores obtained under the specific null hypothesis of independent sources. In order to evaluate the percentile rank for this approach, PAC scores were standardized using the estimated mean and standard deviation of the null distribution, and standardized PAC scores instead of ‘raw’ ones were compared to the ground truth. We generated a null distribution by applying Eq. (13) to the sensor time series, epoching the data on the independent component level, randomly permuting the order of the epochs separately for each IC, and projecting the thereby shuffled components back to sensor space by applying Eq. (14). Note that **X** in Eq. (13) now refers to the EEG sensor time series **Q**. From the surrogate sensor time series, we obtained surrogate source-level PAC scores by applying the same data processing steps as used for the original data including LCMV source projection, within region dimensionality reduction using PCA, and between-region PAC estimation using the MI. We repeated this procedure 100 times to obtain a null distribution of 100 MI scores consistent with a null hypothesis of zero between-site interactions. We used this null distribution in two different ways: in the bivariate PAC experiments, we normalized the obtained true MI scores of every region combination by subtracting the mean of the null distribution and dividing by its standard deviation to obtain MI_norm_. In contrast, in the univariate PAC experiments, we used the null distribution to obtain a p-value to assess the false positive rate (see Section 3.3.5). PAC scores corresponding to p-values below *α* = 0.05 were considered significant.

#### 3.3.5 Performance evaluation

Each experiment was carried out 100 times. We use different metrics to evaluate the performance of different PAC estimators in the EEG-BI and EEG-UNI experiments, respectively. In the bivariate PAC experiments, we were interested to quantify each PAC metric’s ability to detect the simulated ground-truth between-site PAC interactions. In the univariate PAC experiments, no between-site PAC was present. Here we were interested to quantify how prone PAC metrics are to still signal false positive between-site interactions. Please note that in both types of experiments, we only studied across-region PAC detection and do not evaluate the PAC metrics’ abilities to detect univariate PAC within regions.

##### Percentile rank

In the presence of bivariate simulated PAC, we used the percentile rank (PR) to determine how accurately different PAC approaches discriminate regions pairs with ground-truth bivariate PAC from other region pairs. Every region–region combination was assigned a single real-valued FC score from every PAC estimation pipeline. In order to assess a pipeline’s success, we first sorted all FC scores in descending order to obtain the rank vector 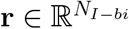, where *N*_*I*−*bi*_ ∈ {1, 2, 3, 4, 5} is the number of ground-truth interactions, and where the jth position of **r** contains the index of the jth ranked connection. The PR was determined using this rank vector:

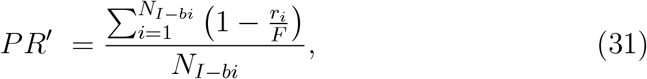

with *F* = *R*^2^ − *R* = 4556 denoting the total number of PAC scores. The *PR*^′^ was then normalized using the perfect-skill PR, *PR*_ps_, and the no-skill PR, *PR*_ns_, and therefore takes values between 0 and 1:

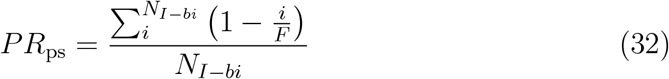

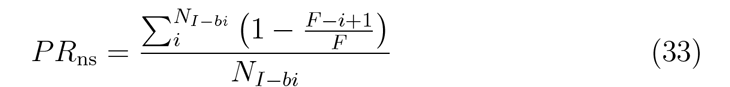

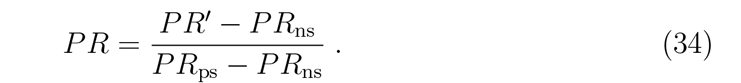

We report all PR values rounded to the second decimal.

##### False positive rate

To define the global FPR, we divided the number of significant PAC interactions (across all region combinations) by the total number of PAC scores *F* :

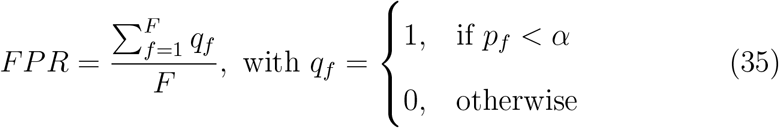

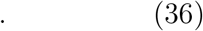

We evaluated the FPR separately for regions that are adjacent to the seed regions with the ground-truth univariate PAC signal, and for regions that are not adjacent to the seed regions. Further, we evaluated the two directions of PAC interactions separately. That is, we calculated the FPR both for PAC between the phase of the seed regions and the amplitude of the neighboring regions, and vice versa.

## 4 Results

### 4.1 Baseline experiment on univariate PAC detection

The baseline experiment assessed the performance of different PAC metrics and pipelines to detect univariate PAC. With this, we wanted to ensure that potential differences in the detection of genuine between-site PAC are not just due to different sensitivities to PAC per se. Figure 4 shows the result of the baseline experiment. We see that all metrics perform similarly well in detecting true within-channel PAC for our chosen filter settings and epoch lengths, with the bispectrum performing slightly worse and the MI by Tort et al. (2010) performing slightly better at -12 dB SNR. To avoid unnecessary high computational cost, we focused on the original MI metric by Canolty et al. (2006) in the following experiments and are not reporting results for the variants proposed by Tort et al. (2010) and Özkurt and Schnitzler (2011).

**Figure 4:**
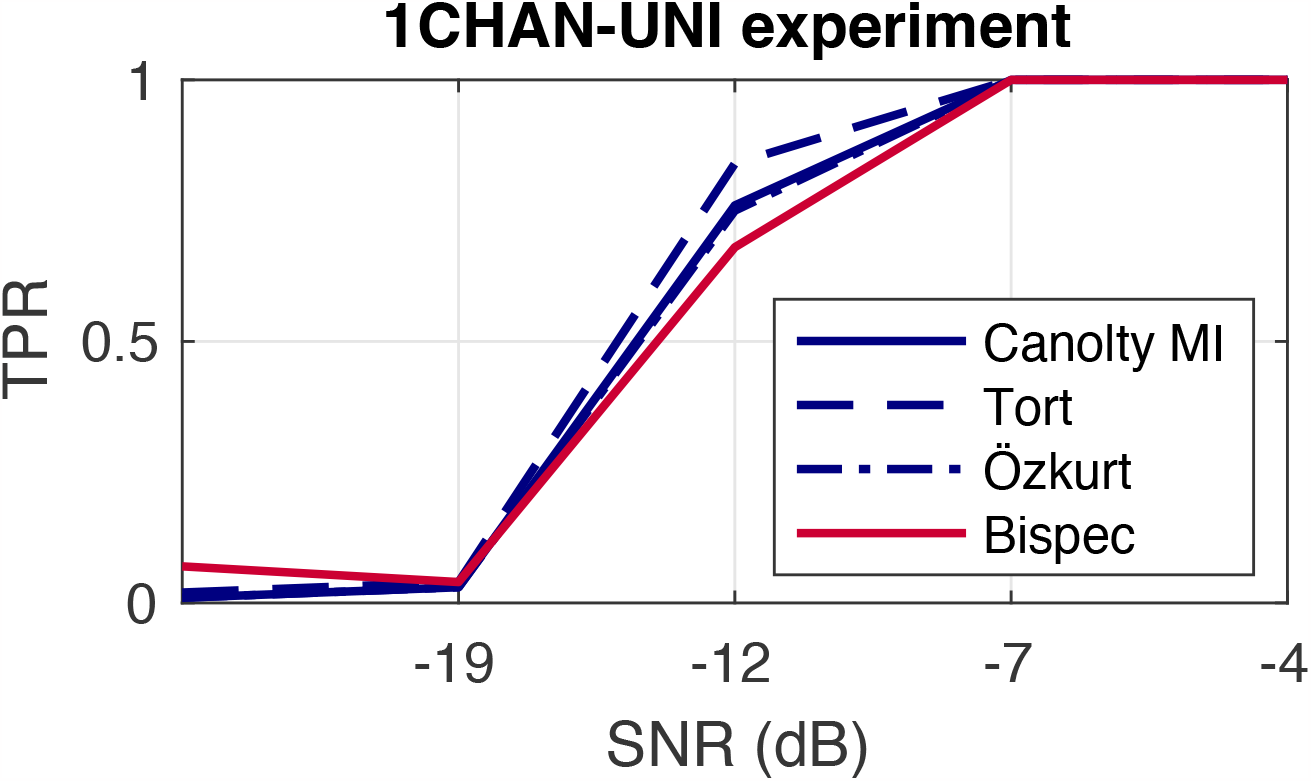
Result of the baseline (1CHAN-UNI) experiment on univariate PAC detection. Sensitivity of mean vector length modulation index Canolty et al. (2006), the two extensions of Tort et al. (2010) and Özkurt and Schnitzler (2011), and bispectrum for the detection of phase-amplitude coupling (PAC) as a function of the signal-to-noise ratio (SNR). PAC is simulated and measured between the slow and fast oscillation within a single channel.

### 4.2 Two-channel experiments

We conducted two types of two-channel experiments: the first type involved true bivariate PAC between the two source channels (2CHAN-BI-IND and 2CHAN-BI-MIX) and the other type involved two source channels containing two independent univariate PAC times series 2CHAN-UNI-IND and 2CHAN-UNI-MIX). The ability to correctly detect the presence of PAC was tested for the modulation index (MI), MI calculated on orthogonalized channels (MI+ORTH), MI tested against surrogate data (MI+ICSURR), the bispectrum (BISPEC), and the anti-symmetrized bispectrum (ASB-PAC).

In Figure 5, we observe that all PAC measures perform well in terms of correctly detecting true bivariate PAC between the two channels. This is also true for random mixtures of the same two channels.

**Figure 5:**
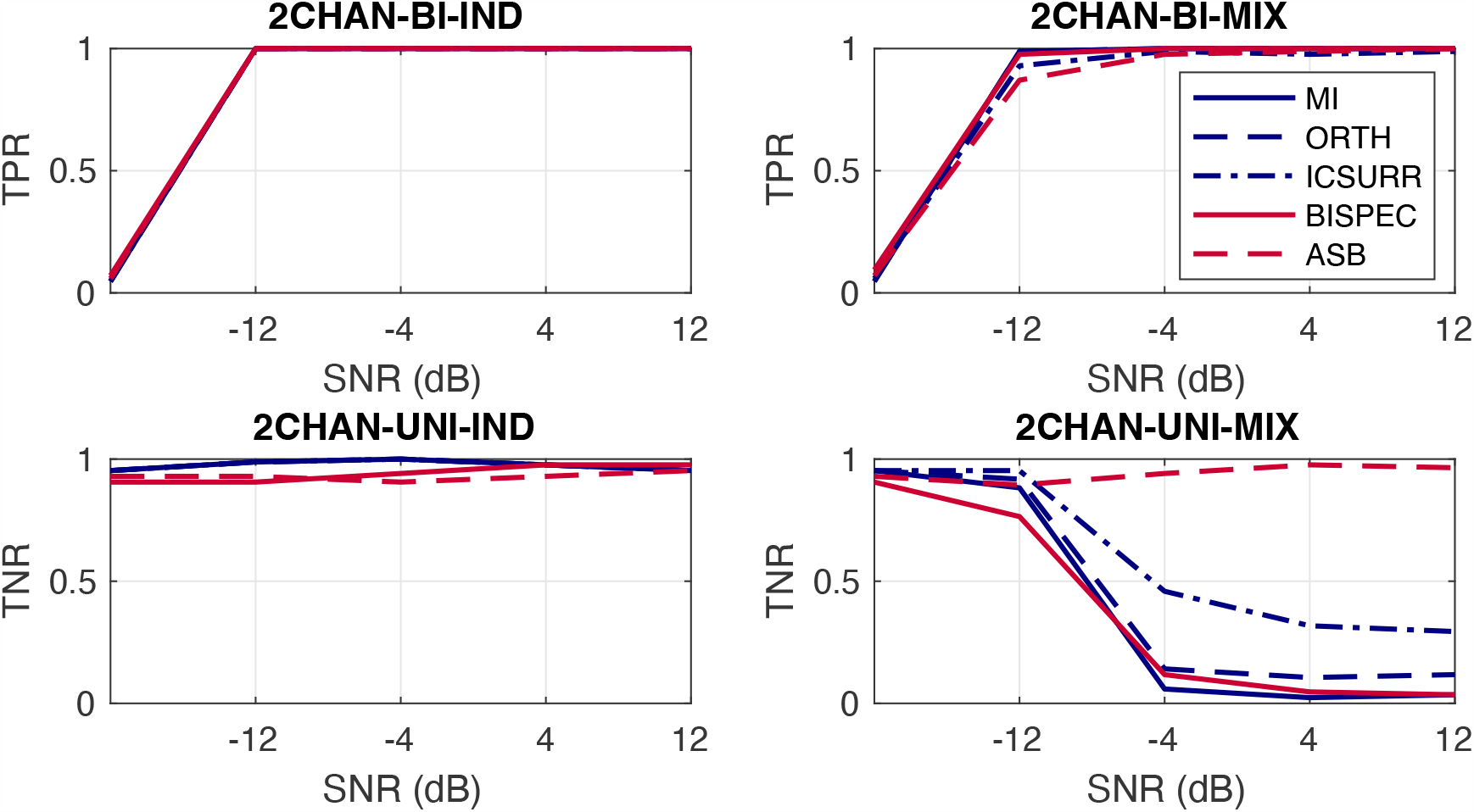
Performance of PAC metrics in two-channel experiments. Top: Simulation of two underlying signals with bivariate PAC without (left) and with (right) additional linear mixing; that is, presence of genuine across-site PAC. Bottom: Simulation of two underlying signals with univariate PAC without (left) and with (right) additional linear mixing; that is absence of genuine between-site PAC. TPR = true positive rate, TNR = true negative rate.

As expected, all PAC metrics avoid falsely detecting PAC almost perfectly in the univariate PAC experiment when there is no signal mixing. However, in case of signal mixing, most metrics indicate PAC in sufficiently high SNR environments. This could be erroneously interpreted as genuine between-channel PAC in practice, even though the PAC interaction stems from a single underlying source. Within the MI-based methods, MI+ICSURR performs best with a TNR of 0.30 for an SNR_t_ of 12 dB (TNR MI = 0.04, TNR BISPEC = 0.04, TNR ORTHO = 0.12 at 12 dB). The only metric that is largely unaffected by signal mixing is the ASB-PAC, which shows a TNR of at least 0.96 throughout all SNRs.

### 4.3 Whole-brain experiments

To assess the ability of the studied pipelines to distinguish between-from within-site PAC under realistic source mixing, we conducted additional experiments on simulated whole-brain EEG data using a realistic volume conductor model.

#### 4.3.1 EEG-BI: detection of true between-site PAC

##### Experiment EEG-BI

In Experiment EEG-BI, we evaluated the performance of five different PAC metrics in distinguishing genuine ground-truth between-region PAC interactions from within-region PAC as well as the absence of any PAC interactions. Figure 6 shows the results of Experiment EEG-BI. We again see that the bispectral metrics perform slightly better than the MI. Importantly, the ASB-PAC does not perform worse than the bispectrum without anti-symmetrization. However, both MI extensions, MI+ORTHO and overall ICSURR, perform worse than the original MI. We discuss this result in Section 7. In the following experiments with bivariate ground-truth PAC, we focus on the MI and the ASB-PAC.

**Figure 6:**
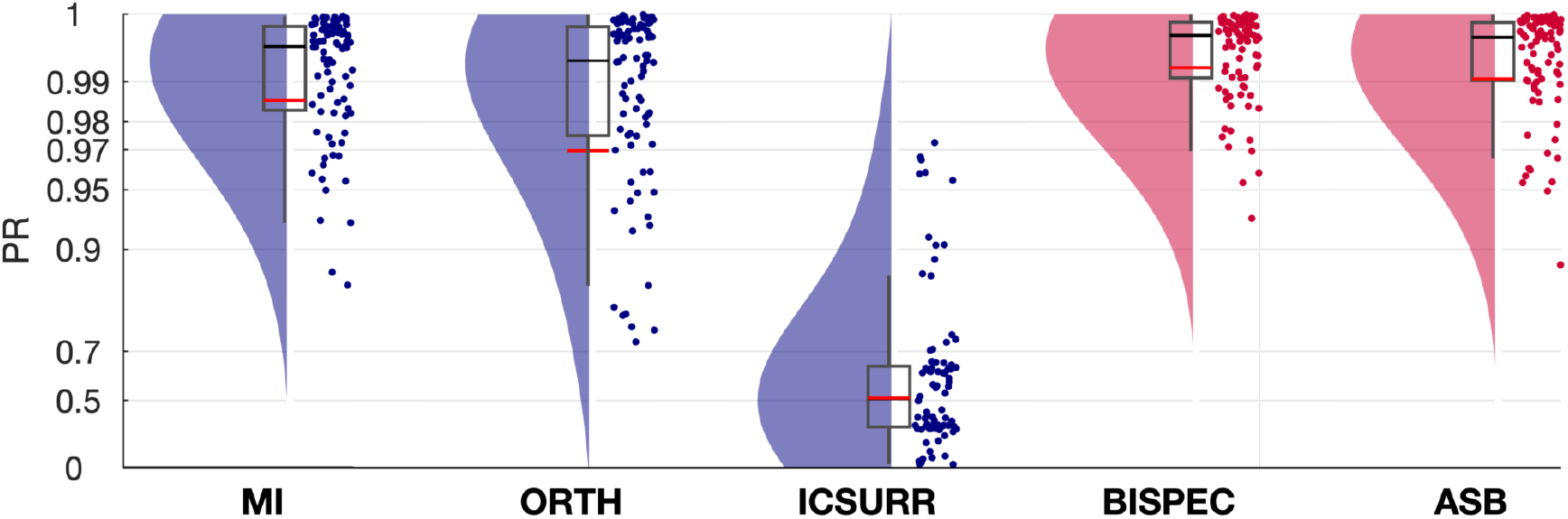
Comparison of different phase–amplitude coupling (PAC) metrics in a whole-brain EEG experiment (Experiment EEG-BI). The dataset encompasses three ground-truth PAC interactions. Pink background noise was introduced to the oscillations, and non-interacting sources were represented using random pink noise signals. EEG signal simulation was conducted using the ICBM152 anatomical head template. Ground-truth sources were allocated in 68 regions of the DK atlas and projected to 97 EEG sensors via a forward model. The synthesized data, comprising signal and noise, had a total SNR of 0 dB and were high-pass filtered at 1 Hz. An LCMV beamformer and principal component analysis were applied to the data, yielding region time courses that underwent PAC metric analysis. Performance was evaluated based on the percentile rank (PR) for the detected bivariate PAC interactions. Red and black lines indicate the mean and median percentile rank (PR), respectively. The boxcar marks the 2.5th and 97.5th percentiles. Note the logarithmically spaced Y-Axis.

##### Experiment EEG-BI-SNR: effect of SNR

In Experiment EEG-BI-SNR, we investigated the effect of the SNR_t_ on the across-region PAC detection performance in the whole-brain EEG setting. In Figure 7, we show the percentile rank attained by MI, BISPEC and ASB-PAC for SNR_t_s of -7.4 dB, 0 dB, and 7.4 dB, respectively. As expected, we see that the performance of all metrics decreases for low SNR_t_.

**Figure 7:**
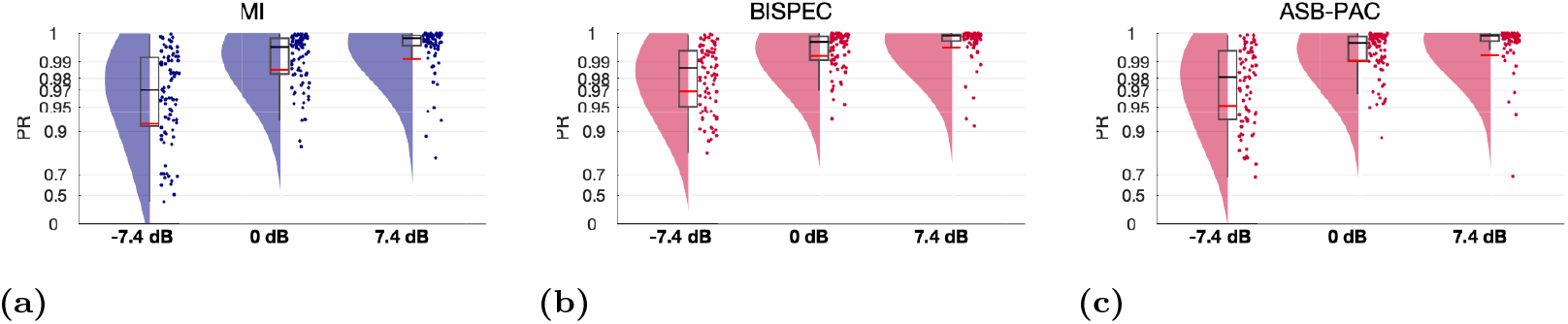
Performance in correctly detecting between-site phase-amplitude coupling as a function of signal-to-noise ratio (Experiment EEG-BI-SNR). Red and black lines indicate the mean and median, respectively. The boxcar marks the 2.5th and 97.5th percentile. Note the logarithmically spaced Y-Axis. It can be seen that the ability to detect true between-site PAC deteriorates in the presence of stronger non-interacting noise signals. MI: Modulation Index, BISPEC: Bispectrum, ASB: Anti-symmetrized bispectrum.

##### Experiment EEG-BI-INT

How does the complexity of interaction patterns affect between-site PAC detection performance? To investigate this, we varied the number of PAC interactions in the ground-truth data in Experiment EEG-BI-INT. In Figure 8, we see that MI, the conventional bispectrum, and ASB-PAC perform worse for many ground-truth interactions than for few interactions.

**Figure 8:**
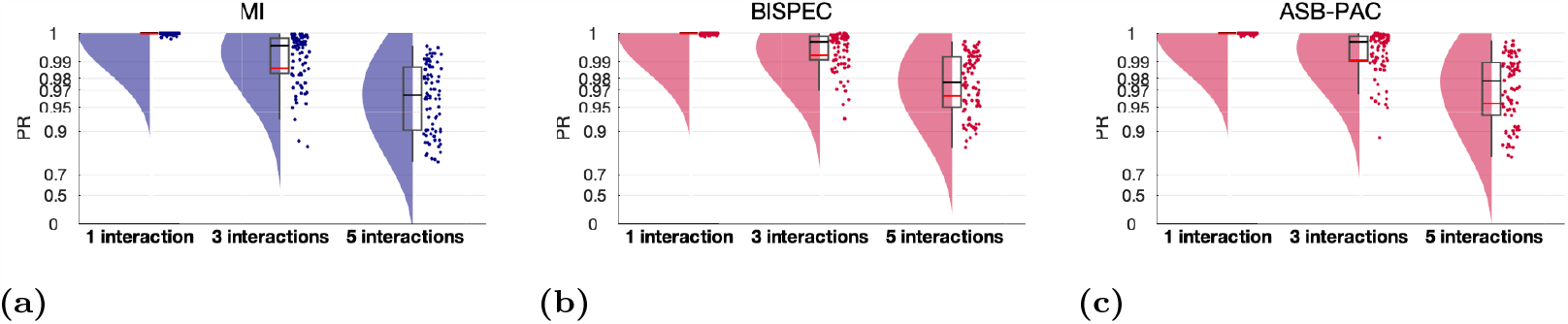
The performance of correctly detecting between-site phase–amplitude coupling depends on the the number of ground-truth PAC interactions (Experiment EEG-BI-INT). Red and black lines indicate the mean and median, respectively. The boxcar marks the 2.5th and 97.5th percentile. Note the logarithmically spaced Y-Axis. MI: Modulation Index, BISPEC: Bispectrum, ASB: Anti-symmetrized bispectrum.

Note that, despite using a normalized version of the PR metric, it is not possible to objectively compare PR scores obtained for different numbers of true interactions. While PR distributions are nearly uniform when only one interaction is simulated, they tend to become more concentrated around 0.5 resembling Gaussian and super-Gaussian distributions with increasing kurtosis for increasing numbers of interactions (Pellegrini et al., 2023).

#### 4.3.2 EEG-UNI: univariate within-site PAC

In Experiments EEG-BI, we observed that bispectral methods perform equally as or superior to the MI at detecting ground-truth bivariate between-region PAC. In Experiment EEG-UNI, we assess the same metrics’ ability to reject spurious between-region PAC in the univariate PAC setting within the whole-brain EEG experimental setting.

Two directions of spurious PAC are conceivable: First, there may be PAC between the phase of the seed region, containing the univariate within-region PAC, and the amplitude of another region. Second, there may be spurious PAC between the amplitude of the seed region and the phase of other regions. It may be that the effect of signal mixing depends on the direction of the coupling. Therefore, we here show the results for both directions.

Further, we distinguish between spurious interactions from the seed region to neighbors and to non-neighbors, since we expect a more extreme impairment for interactions between regions that lie close together and are therefore more affected by source leakage. In Figure 9, we show all 2x2 combinations of these parameters (seed = phase vs. seed = amplitude, neighbors vs. non-neighbors).

**Figure 9:**
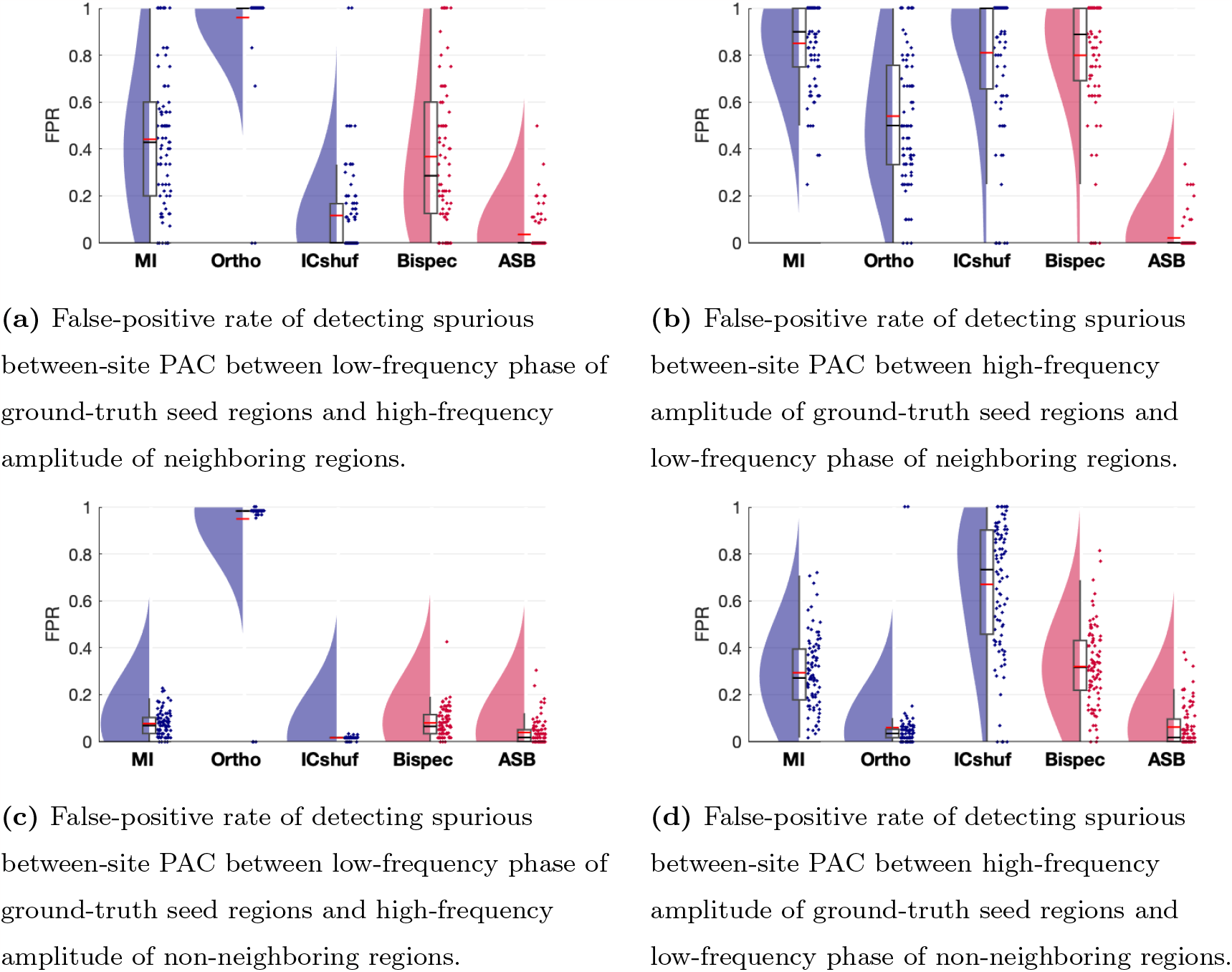
Comparison of different phase–amplitude coupling (PAC) metrics in their ability to avoid detecting spurious between-site PAC (Experiment EEG-UNI). (a/b) Between seed regions and neighboring regions. (c/d) Between seed regions and non-neighboring regions. Red and black lines indicate the mean and median percentile rank (PR), respectively. The boxcar marks the 2.5th and 97.5th percentile.

##### Experiment EEG-UNI

In Experiment EEG-UNI, we compared the different PAC metrics for the default experimental setting. In Figure 9a, we show the FPR associated with the spurious detection of between-site coupling between the phase of the ground-truth seed region containing univariate PAC and the amplitude of the neighboring regions. We observe that the ASB-PAC detects by far the fewest false positives. The MI and the bispectrum without anti-symmetrization cannot eliminate the source leakage effect and detects some false positives. While the ICSURR method seems to improve the MI, orthogonalization results in a very high FPR.

Figure 9b shows the FPR for measuring spurious PAC between the amplitude of the ground-truth seed region containing univariate PAC and the phase of neighboring regions. We see a similar pattern as in Figure 9a. However, both the MI and overall the bispectrum without anti-symmetrization perform even worse. The ASB-PAC detects by far the fewest false positives also in this case.

As expected, these effects are far less pronounced for interactions between the ground-truth seed regions and non-neighboring regions (Figures 9c and 9d). This indicates that the observed differences between the robust ASB-PAC, the non-robust MI and the bispectrum without anti-symmetrization indeed arise from the source leakage that is most pronounced in regions that are adjacent to each other.

##### Experiment EEG-UNI-SNR

In Experiment EEG-UNI-SNR, we investigated how the FPR depends on the SNR, asking:is the ability to reject spurious between-region PAC compromised in low-SNR settings?

In Figure 10, we show the FPR for MI, BISPEC, and ASB-PAC for SNRs of 0 dB, 7.4 dB, and 12 dB. We see that, while the FPR is slightly elevated at 0 dB in case of the MI and bispectrum, the high specificity of the ASB is not compromised at low SNR settings.

**Figure 10:**
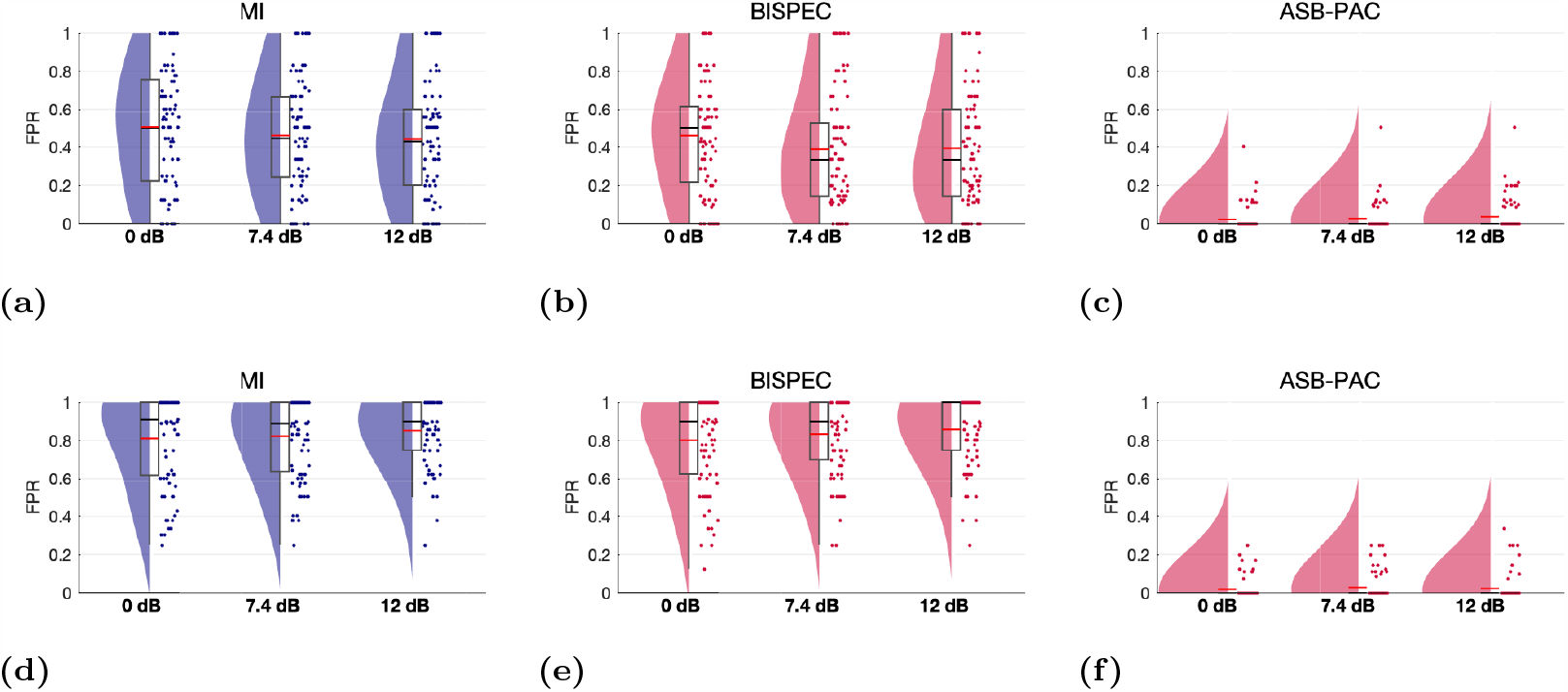
Ability of different PAC estimation pipelines to reject spurious between-site PAC in different signal-to-noise (SNR) settings (Experiment EEG-UNI-SNR). (a) to (c): Rate of false-positive detections of between-site PAC detections between the phase of low-frequency signals at ground-truth seed regions and the amplitude of high-frequency signals at neighboring regions. (d) to (f): Rate of false-positive between-site PAC detections between the amplitude of low-frequency signals at ground-truth seed regions and the phase of high-frequency signals at neighboring regions. Red and black lines indicate the mean and median, respectively. The boxcar marks the 2.5th and 97.5th percentile. MI: Modulation Index, BISPEC: Bispectrum, ASB: Anti-symmetrized bispectrum.

##### Experiment EEG-UNI-INT

In Experiment EEG-UNI-INT, we compared the ability to avoid detecting spurious between-region PAC for different numbers of ground-truth univariate within-site interactions. In Figure 11, we see that the performance of the MI and the bispectrum is slightly compromised for multi-interaction settings. Conversely, the high performance of ASB-PAC does not depend on the number of ground-truth interactions.

**Figure 11:**
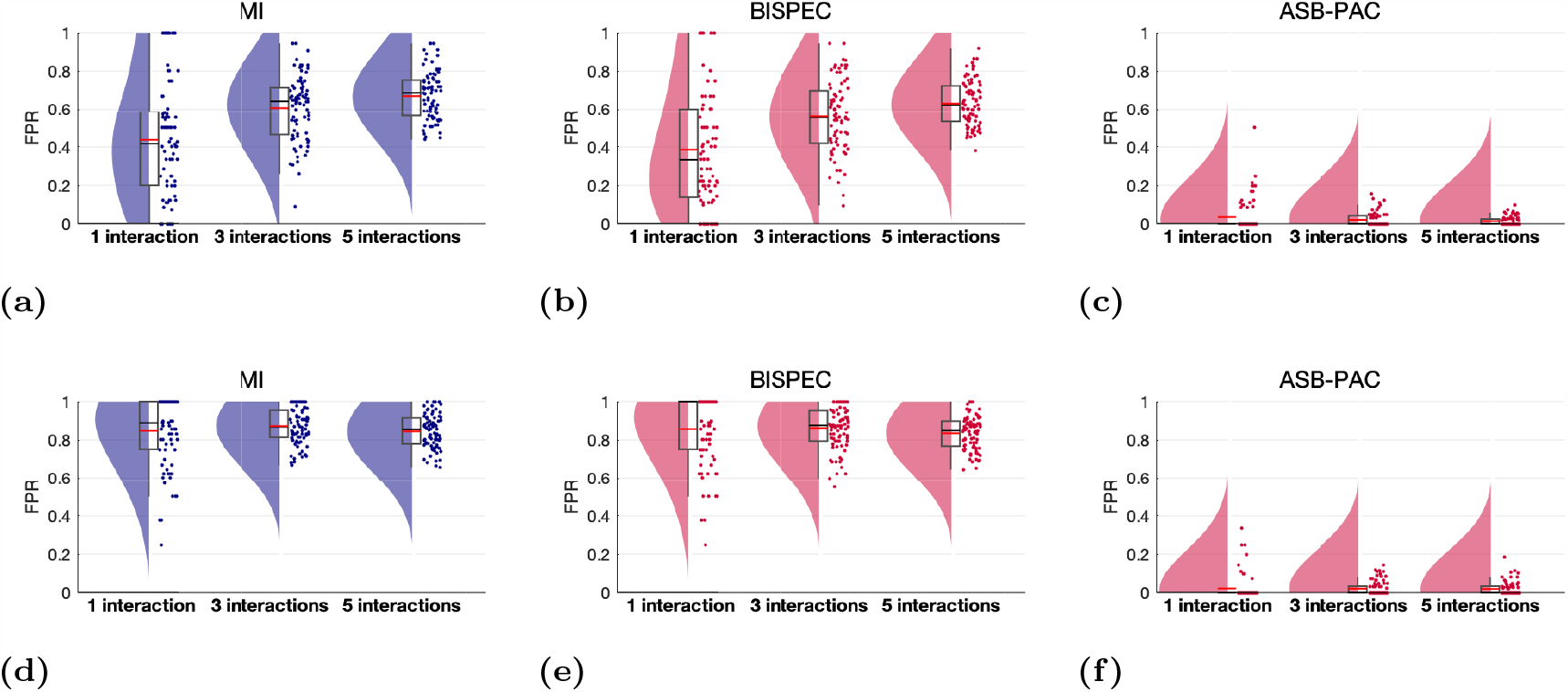
Ability of different PAC estimation pipelines to reject spurious between-site PAC for different number of ground-truth univariate PAC seed regions (Experiment EEG-UNI-INT). (a) to (c): Rate of false-positive detections of between-site PAC between the phase of low-frequency signals at ground-truth seed regions and the amplitude of high-frequency signals at neighboring regions. (d) to (f): Rate of false-positive detections of between-site PAC between the amplitude of low-frequency signals at ground-truth seed regions and the phase of high-frequency signals at neighboring regions. Red and black lines indicate the mean and median, respectively. The boxcar marks the 2.5th and 97.5th percentile. MI: Modulation Index, BISPEC: Bispectrum, ASB: Anti-symmetrized bispectrum.

## 5. Exploratory analysis of PAC in motor imagery

To demonstrate the utility of the ASB-PAC in real EEG data, we present an exploratory analysis of PAC in motor imagery. The Berlin arm of the VitalBCI study (Blankertz et al., 2010; Sannelli et al., 2019) involved 39 participants who took part in an experiment where, among other tasks, they imagined performing a movement with either their left or right hand (referred to as the Motor Imagery Calibration set; MI-Cb 1-3).

Motor imagery (MI) is a mental simulation of movement (Neuper et al., 2005). It has been shown that it induces neural activity in primary sensory and motor areas that is similar to activity incudes by real movements and that can be measured by EEG (Scherer and Vidaurre, 2018). Movement-related EEG responses include event-related potentials (ERPs, e.g., Jongsma et al., 2013) and event-related desynchronization or synchronization (ERD/S Scherer and Vidaurre, 2018) of specific oscillations. For example, it has been shown that motor imagery of hand movements result in an ERD in the *μ* and *β* frequency band (Scherer and Vidaurre, 2018). Motor imagery may also be reflected in functional connectivity patterns, as several studies have shown (Vidaurre et al., 2020; Pellegrini et al., 2023).

It is conceivable that also PAC may emerge during motor imagery, reflecting processes of cognitive integration and communication during the planning of the movement. Technically, bivariate, between-site, PAC may also arise from the co-occurence of both ERP and ERD/S phenomena as a response to the presentation of a stimulus. On the other hand, univariate, within-site, PAC could emerge as an ubiquitous phenomenon that reflects non-linear processes in the data generation, which manifest in specific non-sinusoidal wave shapes of brain oscillations and non-Gaussian distributions. Such univariate PAC coupling can also occur due to non-zero mean of oscillations where for instance the amplitude modulation of alpha oscillations would mimic PAC (Studenova et al., 2022). Thus, it is an interesting question whether and between which sites PAC occurs during motor imagery. Furthermore, if PAC is observed, one may ask whether the observed phenomenon has an interpretation as an interaction between distinct brain regions (between-region PAC) or could be explained by single individual PAC sources (within-region PAC). Conversely, if PAC is observed due to a co-occurrence of ERP and ERD/S, it is of interest whether these are potentially elicited by the same sources or provably come from distinct generators.

In every trial of the VitalBCI experiment, a visual stimulus displaying a fixation cross accompanied by an arrow indicating the task (i.e., left or right motor imagery) was shown. Four seconds later, the stimulus disappeared, and the screen remained black for 2 seconds. Subjects were asked to perform motor imagery of the respective hand for as long as the arrow was present on the screen. Each subject completed 75 left and 75 right MI trials. EEG data were recorded using a 119-channel whole-head EEG system with a sampling rate of 1000 Hz (see Blankertz et al., 2010; Sannelli et al., 2019, for further details).

For the analysis, we used the ROIconnect plugin^3^ (Pellegrini et al., 2023) in combination with EEGLAB. We used a standard set of 90 EEG channels covering the whole head. Further, we selected 26 subjects for our analyses.

The subject inclusion criteria were based on previous studies that demonstrated successful classification between left and right motor imagery conditions using machine learning methods (“Category I” in Sannelli et al., 2019). Prior to the analysis, the data underwent several preprocessing steps, including filtering (1 Hz high-pass filter, 48–52 Hz notch filter, and 45 Hz low-pass filter, all implemented using zero-phase forward and reverse second-order digital Butterworth filters), as well as subsampling to 100 Hz. Artifactual channels were identified through visual inspection of the power spectrum and the topographical distribution of alpha power. On average, 1.19 channels (ranging from zero to five per participant) were found to be artifactual and were subsequently interpolated using spherical scalp spline interpolation. A leadfield was computed using the Colin27 5003 Standard-10-5-Cap339 template head model (5003 voxels, standard 10-5 channel positions), which is a preexisting component of the EEGLAB toolbox. Subsequently, the data were cut into segments ranging from 1 to 3 seconds after the start of each stimulus presentation and separated into left and right motor imagery trials. For further analysis, only the right-hand motor imagery trials were used. To project sensor data to source level, we used an LCMV filter which was constructed on the same sensor data. Finally, we aggregated the multivariate source time series within regions using PCA and selected the strongest PC per region (see Pellegrini et al., 2023).

Previous research suggests that functional connectivity is modulated in motor imagery (Vidaurre et al., 2020; Pellegrini et al., 2023) with especially the sensori- and motor cortices being involved (Scherer and Vidaurre, 2018). However, PAC between the different involved areas has never been investigated before with robust metrics. In this study, we selected the left and right pre- and post-central cortices as regions of interest, as this is where the sensori- and motor areas are located.

In the following analysis, we addressed the following questions:

1. Is motor imagery characterized by within-site PAC during in the sensori- and motor cortices?
2. Is motor imagery also associated with genuine between-site PAC between sensori- and motor cortices, indicating a role of PAC as a mechanism of distant brain communication in MI?
3. If PAC as detected by the conventional bispectrum is observed, does it vanish when using a robust PAC metric, i.e., the ASB-PAC? We hypothesized that the differences between the between-site PAC estimates obtained by the uncorrected bispectrum and the ASB-PAC are more pronounced for regions that lie close together, in contrast to regions that lie on different hemispheres.

To this end, we estimated PAC in three ways:

- Within-region PAC measured with the conventional bispectrum (BIS-PEC).
- Between-region PAC measured with the conventional bispectrum (BIS-PEC).
- Between-region PAC measured with ASB-PAC.

Since previous research found that motor imagery is mostly reflected in the *μ* and *β* bands, we calculated bispectra between slow oscillations of 1 to 12 Hz, and fast oscillations between 1 and 50 Hz (both with 1 Hz resolution). Since Fourier coefficients can only be evaluated up to the Nyquist frequency in a meaningful way, and the third term of the bispectrum includes the sum of the slow and fast oscillation frequencies, our analyses had to be restricted to frequency combinations whose sum did not exceed the Nyquist frequency. To ensure that the fast oscillation frequency carrying the amplitude envelope is significantly higher than the slow oscillation frequency carrying the phase, and to avoid strong confounds by interactions of the slow oscillation with its second and third harmonic, we constrained our analyses to fast oscillation frequencies whose frequency was at least three times higher than the frequency of the corresponding slow oscillation.

To test whether the observed PAC scores are significantly different from chance levels, we employed the permutation-based statistical approach as in the simulations (see Section 2.4). In brief, for each observed PAC score, we calculated the same bispectral metrics for randomly across epochs permuted Fourier coefficients to obtain samples of a null distribution consistent with the null hypothesis of no PAC interaction (see Eq. 8). We generated 5000 samples of the null distribution and subsequently evaluated for how many of the surrogate samples the estimated PAC value exceeded the PAC value obtained on the original data (see Eq. 9). To aggregate the resulting p-values over subjects, we employed Stouffer’s method (Dowding and Haufe, 2018).

Figure 12a shows frequency combinations with significant PAC as estimated with the conventional bispectrum without anti-symmetrization. We observe significant within-as well as between-region PAC between the phase of low-frequent oscillations and the amplitude of beta and gamma oscillations across many frequency combinations and between all ROIs. This leads to the question to what extent the observed between-region effects indeed reflect underlying coupling between distinct brain areas as opposed to non-linear properties of single brain sources.

**Figure 12:**
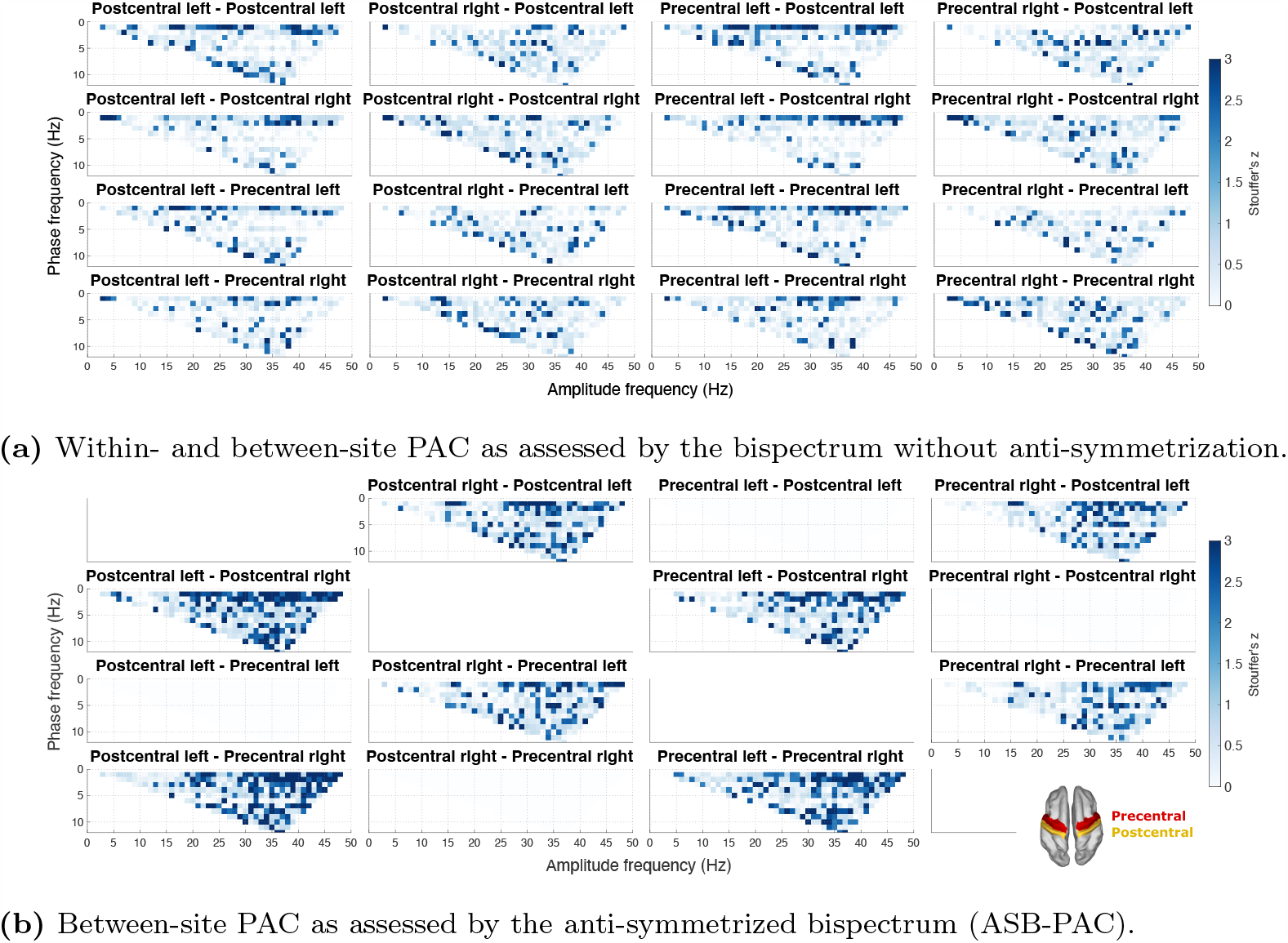
Phase-amplitude coupling (PAC) within and between left and right postcentral and precentral cortices during motor imagery of the right hand. Displayed are z-scores resulting from Stouffer’s method of combining subject-level p-values. In both figures, only statistically significant (p *<* 0.05) effects are drawn opaque. The thumbnail on the bottom right of (b) shows the cortical localization of the two regions of interest.

Figure 12b shows corresponding results for ASB-PAC. We clearly see that the observed PAC between regions in the same hemisphere vanishes when employing the anti-symmetrization. This absence in comparison to Figure 12a suggests that part of the between-region PAC implicated by the original bispectral metric in fact cannot be explained by genuine between-site interactions and may instead results from signal spread between the pre- and post-central cortices of the same hemisphere. In contrast, inter-hemispheric PAC effects were even more pronounced when assessed through ASB-PAC as compared to the conventional bispectrum. This may speak to a favourable cleaning effect of the anti-symmetrization by which signal contaminations attributable to signal spread are effectively removed.

Matlab code of the analyses presented in this section is provided under^4^.

## 6. EEGlab plugin for bispectral PAC estimation

ROIconnect is a freely available open-source EEGLAB plugin that provides a suite of signal processing methods proposed in the literature to estimate FC between regions of interests (ROIs). We first introduced the toolbox in Pellegrini et al. (2023), describing basic features including source reconstruction techniques, dimensionality reduction within regions, region-based power estimation with optional 1/f correction and estimation of region-based 1/f slope, robust inter-regional linear FC estimation, as well as various options to visualize power and FC. All functions can be accessed through the EEGLAB GUI or the command line. As an EEGLAB plugin, ROIconnect has access to core EEGLAB functions for importing and preprocessing EEG data as well as for calculating the leadfield and the source model. The toolbox is available on GitHub under^5^ and is installable through the EEGLAB GUI extension manager. In the following section, we describe advanced features and main updates to the toolbox resulting from the outcomes of this study.

### 6.1 Phase-amplitude coupling

Our results suggest that the bispectrum is a suitable method to detect PAC. Further, we observed that the ASB robustly detects between-site PAC in the presence of volume conduction considerably better than the conventional bispectrum or methods based on the Modulation Index. To this end, we implemented functions for robust estimation of (within- and between-region) PAC based on the bispectrum, and for between-region PAC based on the ASB.

The `pop_roi_connect` function of ROIconnect allows users to estimate PAC between two frequencies or frequency bands, generating bispectral estimates for the required region-region-frequency-frequency combination. The number of ROIs can be predefined by the user. Beyond PAC, this function can also be utilized for other functional connectivity metrics calculations.

### 6.2 Statistics

A statistics mode is provided for assessing the statistical significance of the estimated FC metrics using a permutation-based approach. For bispectral connectivity metrics, we implemented the method used in this study, where a surrogate distribution sample is obtained by randomly permuting the epochs of a channel. The cross-bispectrum is then computed on the shuffled time series.

Statistical analysis of FC metrics is available through `pop roi connect` by activating the statistics mode. Additionally, the resulting p-values can be visualized as cortical surface topographies for a selected frequency or frequency band.

## 7. Discussion

In this study, we addressed potential limitations of existing PAC metrics, such as the MI and bispectrum, in assessing genuine between-site PAC and their vulnerability to signal mixing. We evaluated the effectiveness of three strategies, namely the ASB-PAC, MI with orthogonalization, and the IC surrogate approach, to enhance the robustness of the conventional PAC metrics. Furthermore, we explored the influence of the SNR and the number of ground-truth interactions on PAC detection accuracy.

We introduced a minimal two-channel experimental setting, which allowed us to quantify and compare the performance of various PAC metrics in correctly distinguishing the presence of ground-truth between-channel PAC from univariate PAC observed in the same channel pairs but originating from single sources. We identified ASB-PAC to be the only PAC metric that is able to consistently reject spurious between-channel PAC in a setting with two independent but mixed univariate PAC time series, while also being able to detect genuine ground-truth bivariate (between-site) interactions with high sensitivity. In a more complex whole-brain EEG-like experimental setup, we confirmed the excellent performance of ASB-PAC in both cases: it robustly detected ground-truth inter-regional PAC in low to moderate noise settings. Conversely, in the exclusive presence of univariate PAC sources, ASB-PAC attained a low false-positive rate, whereas other metrics were characterized by the detection of many spurious interactions.

### Spurious between-site PAC is a result of local source leakage

False-positive between-site PAC was more frequently detected between neighboring regions in comparison to non-neighboring regions. This is an expected result since so-called source leakage occurs predominantly between regions that lie close together. This can also be seen in Figure S1, which shows that there is a higher correlation between the time courses of a seed region and its neighbors compared to non-adjacent regions. This is then also reflected in higher but spurious PAC (Figure S2).

We found more spurious PAC between the amplitude of the fast oscillation in the seed PAC region and the phase of the slow oscillation in the neighboring regions than vice versa. This can be explained by the comparably higher power or SNR of the univariate PAC signal at low frequencies due to the applied 1/f scaling (see Section 3.3.1). The higher power leads to more source leakage for the slow oscillation than for the fast oscillation. Therefore, false-positive between-region PAC is more likely to be observed between the phase of this leaked slow oscillation and the fast oscillation-amplitude measured in the seed region. We investigated this in a single whole-brain experiment (univariate PAC, all parameters set to default), where we assessed the correlation between the time courses of the slow and fast oscillation of the seed region and the corresponding slow and fast oscillation of the neighboring regions, respectively (see Figure S1). We found a mean absolute Pearson correlation of *r* = 0.42 for the slow oscillation, but a correlation of only *r* = 0.22 for the fast oscillation, which shows that the slow oscillation leaked more from the seed region into the neighboring regions than the fast oscillation due to its higher SNR_t_.

### Physiological interpretation of ASB-PAC

Note that, while the bispectral index 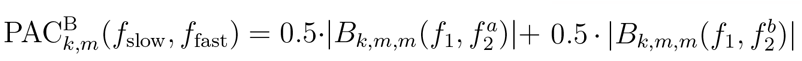 is a valid characterization of PAC, this is not necessarily the case for the difference measure 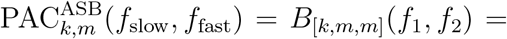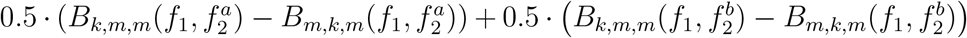 meaning that there might in general be no real-world PAC signal whose bis-pectrum is PAC^ASB^. Specifically, the correction terms 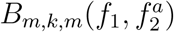 and 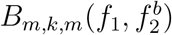 correspond to interactions between a signal *x*_*m*_ at frequencies *f*_1_ and 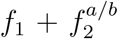 with another signal *x*_*k*_ at frequencies 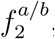, respectively, which, as such, are not PAC interactions. One may thus wonder what it means to subtract *B*_*m,k,m*_ from *B*_*k,m,m*_, and whether the subtraction could potentially lead to the removal or cancellation of neurophysiological information of potential interest. Here we argue that this is unlikely, as nonlinear interactions between 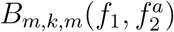 and 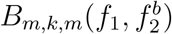 as described above are not expected to occur regularly in the brain. On the other hand, such interactions provably emerge as a result of source mixing in the presence of a source with within-site PAC. In such cases, it is beneficial to remove the leakage artifact picked up by 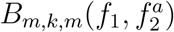 and 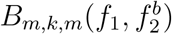, whereas in cases without substantial leakage, both terms are expected to be negligible and thus their removal is not expected to influence the ASB-PAC. This consideration is confirmed by our numerical experiments, where the sensitivity (as measured by the PR) of the ASB-PAC in the bivariate PAC setting is comparable to that of the original bispectrum while a consistently low FPR is attained in the presence of solely univariate ground truth PAC interactions throughout all mixing proportions and SNRs.

A situation in which what we call genuine between-site PAC, as characterized by ASB-PAC, still has ambiguous interpretations is when one signal with univariate PAC is reflected in two channels with different constant time delays. The resulting between-channel PAC cannot be explained by linear mixing as a volume conduction or source leakage artifact. Yet, one may argue that the simplest and thus more likely mechanism of information transfer in this setting is just a mere delayed forwarding of the broad-band signal than an intricate phase-amplitude coupling between distinct frequency bands. To test for such simpler hypotheses, it can be advisable to additionally test for stable non-zero phase delays between channels either in the low frequency band, the high frequency band, or both. This can be achieved using established metrics based on the cross-or bispectrum (Nolte et al., 2004; Haufe et al., 2013; Winkler et al., 2016).

### Orthogonalization and ICA-based surrogates cannot distinguish within-from between-site PAC

Both orthogonalization (Colclough et al., 2015; Hipp et al., 2012) and surrogate data approaches perturbing the data at the level of independent components (Shahbazi et al., 2010) have been proposed as general purpose strategies to remove artifacts of volume conduction from M/EEG data and subsequent functional connectivity analyses. Thus, it may seem worthwhile to assess the ability of these approaches to distinguish genuine between-site PAC from PAC that is measured between two different sites but is inherently caused by within-site PAC of a single sources whose signal leaks into both sites. Our experimental results, however, demonstrate that both the orthogonalization approach and the ICA-based surrogates do not improve the performance of the MI in terms of detecting the presence of across-region PAC. Orthogonalization even worsens the performance of the original MI metric in the whole-brain experiments. The use of ICA based surrogates seems to provide an improvement in certain simple setups such as the two-channel experiments with source mixing but not in more complex scenarios with larger numbers of channels.

Here we provide arguments for the suboptimal behavior of both methods for the studied problem. Both approaches decompose the sensor data into components that are orthogonal and (in case of ICA-based surrogates) “maximally” statistically independent from another. Here statistical independence is typically achieved by maximizing non-Gaussianity based on the argument that mixtures of independent signals (e.g. due to volume conduction) are closer to be Gaussian distributed than the unmixed signals. The idea of the ICA-based surrogate data approach is to remove any residual functional connectivity at the component level by randomly shifting component time course relative to each other. When projecting the shifted components back to sensor space, surrogate data complying with the null hypothesis of zero source-level FC are hoped to be obtained, which can be tested against. However, this reasoning only applies to FC metrics that are strictly bi-or multivariate. Cross-frequency interactions such as PAC, which can emerge within single univariate time series, are likely to be retained in single independent components, and are thus preserved even under random shifts of the components relative to each other. We can easily understand that exactly that happens for univariate PAC sources, whose individual slow and fast-oscillation spectral components are Gaussian distributed by construction, but the summation of which is (by virtue of the higher-order PAC interaction) super-Gaussian. Thus, ICA algorithms aiming to maximize non-Gaussianity have a strong incentive to retain univariate PAC signals in single components rather than splitting them into slow- and fast-oscillation parts. We confirmed this behavior by investigating power spectra of the ICs, calculated from sensor-level activity in the EEG-UNI experiment in one simulation run. We found that a large number of ICs contained both the low-frequency peak and the high-frequency peak with the two side lobes (Figure S3). Consequently, permuting the components does not create an adequate null distribution. Spatio-spectral decomposition (SSD, Nikulin et al., 2011) could be a more suitable approach to overcome this problem. In contrast to ICA, it is designed for maximizing the signal power at specific frequencies while suppressing signal power at other frequencies. The method could be used to force the slow and fast oscillation into two different components, and permutation could then result in a more appropriate null distribution. The proposition of SSD-based surrogates as a novel methodology is, however, beyond the scope of this study. Using ICA decomposition and PAC estimation, Gong et al. (2021) showed that the strength of cortical PAC between the phase of beta oscillations and the amplitude of high-frequency gamma activity was increased in patients with Parkinson’s disease compared to healthy participants. Moreover, PAC also predicted the severity of motor deficits measured with the part 3 of Movement Disorder Society Unified Parkinson’s Disease Rating Scale. Importantly, this prediction was only possible when the phase and amplitude of the corresponding PAC oscillations was estimated with spatially distinct independent components but not for the PAC within the same components. A validation of these findings with the methods presented in this paper, would provide further evidence for the existence of such across-site PAC. This in turn would be important for the introduction of novel methods for the multilocus non-invasive brain stimulation (Koponen et al., 2018) aimed at the disruption of pathological PAC.

The orthogonalization approach of Colclough et al. (2015) aims to remove artifacts of volume conduction by finding the set of time series that are closest to the original sensor time series in the mean-squared error sense, yet are uncorrelated. This and similar approaches are often used with the implicit expectation that the removal of first-order correlations between orthogonalized time series would also destroy higher-order interactions introduced by the same source mixing process and enable the analysis of non-linear interactions such as PAC or amplitude envelope correlations (AEC, Hipp et al., 2012; Colclough et al., 2015) without interference from artifacts of volume conduction. However, this is only guaranteed if the orthogonalization indeed recovers the original unmixed sources, which can be prohibited if the number of underlying sources is larger than the number of sensors and/or if the mixing matrix (in the EEG setting the so-called resolution kernel composed of the leadfield and inverse projection matrix) substantially deviates from diagonality. In addition, slow and fast oscillation spectral components of a PAC interaction are in general linearly uncorrelated. Thus, any orthogonalization algorithm has no incentive to group slow and fast oscillation signal parts into the same component but may just as well assign slow and fast oscillation parts to different components that would be uncorrelated but PAC coupled even if the original PAC interaction was just within site. This would lead to the emergence of spurious between-site PAC even on orthogonalized signals.

These considerations signify that tailored solutions such as antisymmetrized bispectra are needed to distinguish genuine between-from within-site PAC. Similarly, simulations as performed here are advisable to critically assess the aptitude of approaches such as orthogonalization when used to address other novel problems in brain functional connectivity estimation.

### Limitations

In this study we observed that PAC metrics based on bispectra are well suited for detecting ground-truth PAC even in challenging SNR regimes, and that anti-symmetrization can further robustify the estimation in the sense that it prevents the detection of between-site PAC that can be more trivially explained by mixtures of univariate PAC sources. However, we did not investigate the possibility of spurious (within-or between-site) PAC due to other reasons. There are various additional possible pitfalls that warrant further discussion/investigation.

First, while the bispectral indices used here do reflect PAC, they may also be influenced by other types of coupling, like phase–frequency, amplitude–frequency, and amplitude–amplitude interactions (Hyafil et al., 2015; Jirsa and Müller, 2013). These possibilities should be carefully ruled out when investigating PAC with bispectra.

Second, it has been shown that spurious PAC detection may arise from physiological artifacts, like eye movements, muscle activity, or heart beat, which can often be simultaneously observed in multiple channels (Giehl et al., 2021). However, this would not pose a problem for ASB-PAC based PAC estimation, which is invariant to mixtures of non-interacting sources by construction (Chella et al., 2014).

And, third, spurious PAC may arise from a rhythmic non-sinusoidal signal and its higher harmonics (Hyafil, 2017; Giehl et al., 2021; Idaji et al., 2022). Indeed, in a recent resting-state MEG study (Giehl et al., 2021), all within-region PAC could be attributed to the present of higher harmonics or physiological artifacts in the data. To rule out unwanted effects of higher harmonics, a novel method, Harmoni (Idaji et al., 2022), was recently developed. Further research should investigate how Harmoni can be combined with the ASB-PAC approach.

## 8. Conclusion

Between-site PAC can spuriously emerge when a univariate PAC signal is spread to other locations. Therefore, a robust method to disentangle genuine between-site PAC from within-site PAC is needed. In this study, we tested the use of the anti-symmetrized bispectrum to robustly estimate genuine between-site PAC. Previous studies have demonstrated that anti-symmetrized bispectra vanish for mixtures of independent sources. However, the application of ASB-PAC to assess the presence of genuine between-site PAC has not been explored yet. To investigate the performance of different algorithms in detecting PAC in the context of mixed signals, and, thus, in distinguishing between genuine between-site PAC and within-site PAC, we conducted two experiments: one using a minimal two-channel setup and one involving a more complex EEG-like setting that mimicked the generation and reconstructions of underlying EEG sources with forward and inverse modeling techniques. Our findings reveal that the ASB-PAC exhibits superior performance in detecting simulated PAC in the presence of volume conduction, outperforming conventional PAC estimators. Specifically, the ASB-PAC approach demonstrated the highest performance in detecting genuine between-site PAC interactions while detecting the fewest spurious interactions in presence of signal mixing. In light of these results, the use of ASB-PAC-based metrics could significantly enhance the interpretation of future studies investigating PAC as a mechanism of brain communication across macroscopic sites. By effectively addressing the issue of spurious between-site PAC emergence in mixed signal settings, the ASB-PAC approach offers valuable insights into the genuine functional interactions between distinct brain sites, thereby facilitating a more accurate understanding of PAC’s role in brain dynamics and signaling processes.

## Data and code availability

The code for the simulation can be found here: https://github.com/fpellegrini/PAC. The code for the ROIconnect plugin can be found here: https://github.com/sccn/roiconnect. And the code for the minimal real data example here: https://github.com/fpellegrini/MotorImag. Data of the real data example are available upon request.

## Author Contributions

**Franziska Pellegrini**: Methodology, Software, Investigation, Writing – original draft, Visualization. **Tien Dung Nguyen**: Software, Writing – original draft, Writing – review & editing. **Taliana Herrera**: Methodology, Investigation. **Vadim Nikulin**: Writing – review & editing, Supervision. **Guido Nolte**: Methodology, Writing – review & editing, Supervision. **Stefan Haufe**: Conceptualization, Methodology, Validation, Investigation, Resources, Writing – review & editing, Supervision, Project administration, Funding acquisition.

## Acknowledgements

This project was supported by the European Research Council (ERC) under the European Union’s Horizon 2020 research and innovation programme (Grant agreement No. 758985) and through Deutsche Forschungsgemeinschaft (DFG, German Research Foundation) – Project-ID 424778381 – TRR 295(Projects B05 and B02). G.N. was partially funded by the German Research Foundation (DFG, SFB936 Z3 and TRR169, B4). The computations for this work were partly run on the open Neuroscience Gateway cluster (Sivagnanam et al., 2013).

## Footnotes

1 https://github.com/fpellegrini/PAC

2 https://www.nitrc.org/projects/meth/

3 https://github.com/sccn/roiconnect

4 https://github.com/fpellegrini/MotorImag

5 https://github.com/sccn/roiconnect

