## Supplement for "Distinguishing between- from within-site phase-amplitude coupling using antisymmetrized bispectra"

---

---

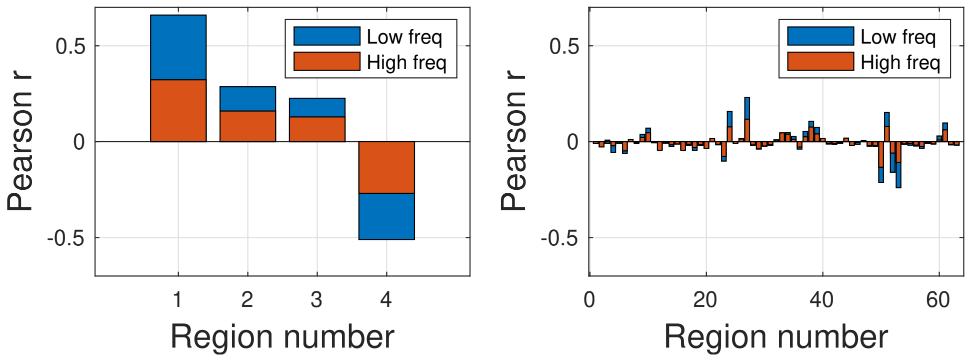


**Figure S1:** Pearson correlation between the reconstructed time courses of a single seed region and its four neighbors (left) compared to the correlation between the seed region and non-adjacent regions (right). Shown data correspond to a single run of the EEG-UNI experiment.

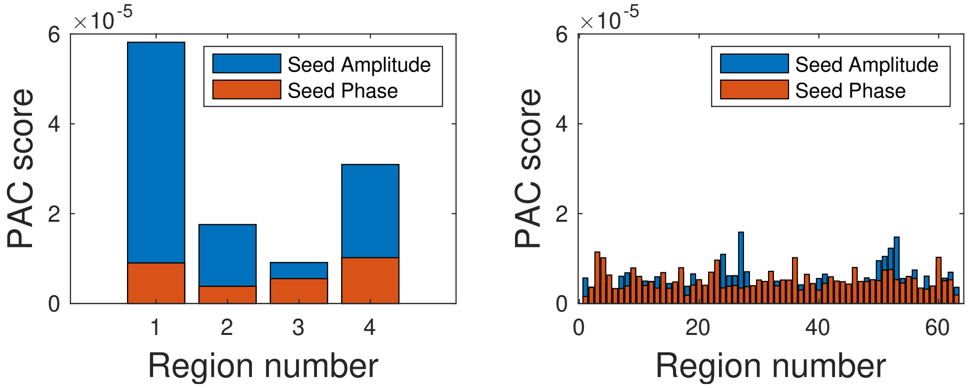


**Figure S2:** Phase-amplitude coupling (PAC) as measured by the bispectrum without anti-symmetrization, between the reconstructed time courses of a single seed region and its four neighbors (left) compared to the PAC between the seed region and non-adjacent regions (right). Shown data correspond to a single run of the EEG-UNI experiment.

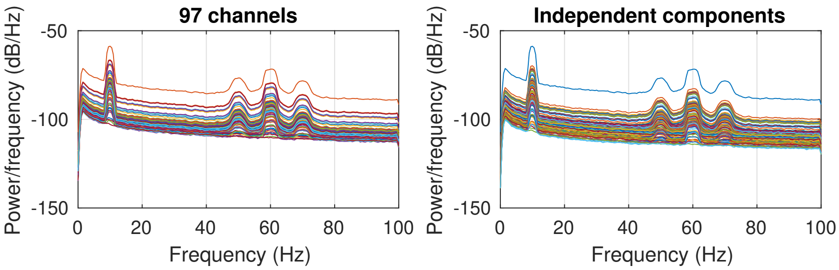


**Figure S3:** Power spectra of the 97 channels (left) and the independent components (right), calculated from sensor-level activity in the EEG-UNI experiment in one simulation run. A large number of ICs contains both the low-frequency peak (10 Hz) and the high-frequency peak (60 Hz) with the two side lobes (50 Hz and 70 Hz)
